# Blood-borne sphingosine 1-phosphate maintains vascular resistance and cardiac function

**DOI:** 10.1101/2025.01.15.633211

**Authors:** Ilaria Del Gaudio, Philippe Bonnin, Emilie Roy-Vessiers, Estelle Robidel, Manuella Garcia, Coralyne Proux, Alexandre Boutigny, Veronique Baudrie, Hoa TT Ha, Ludovic Couty, Nicolo Faedda, Nesrine Mebrek, Théo Morel, Anja Nitzsche, Stéphanie Baron, Olivia Lenoir, Pierre-Louis Tharaux, Maria-Christina Zennaro, Long N. Nguyen, Daniel Henrion, Eric Camerer

**Author notes:** Corresponding authors: Philippe Bonnin, APHP, Hôpital Lariboisière, Service de Physiologie Clinique, 2, rue Ambroise Paré, 75010 Paris, France Eric Camerer, Université Paris Cité, PARCC, INSERM U970, 56 Rue Leblanc, F-75015 Paris, France.

## Abstract

G protein-coupled receptors (GPCRs) are key regulators of cardiovascular function that provide targets for the treatment of cardiovascular disease. Sphingosine 1-phosphate (S1P) is an erythrocyte- and platelet-derived lipid mediator with cognate GPCRs on endothelial cells (EC), vascular smooth muscle cells (VSMC) and cardiomyocytes. S1P circulates in plasma bound to apolipoprotein M (ApoM)-containing high-density lipoproteins (HDL) and to albumin. Circulating S1P levels correlate positively with systolic blood pressure in hypertension and negatively with severity in septic shock and with left ventricular (LV) function in coronary heart disease. In mice, impaired S1P binding to HDL or signaling to EC both trigger hypertension, supporting an essential role for HDL-S1P in supporting endothelial function. The roles of albumin-S1P and myocyte S1PRs in cardiovascular homeostasis remain incompletely defined.

Contrasting isolated HDL-S1P deficiency, we report that non-selective depletion of circulating S1P pools in mice impairs LV contractile function and induces hypotension and resistance to the spontaneous increase in blood pressure with age. Cardiac output was preserved in naïve S1P deficient mice by compensatory LV dilation, but cardiac reserve reduced in a dobutamine stress test. These phenotypes tracked with hematopoietic cell S1P production and were partially or fully reversed by erythrocyte transfusion. Hypotension was accompanied by reduced peripheral resistance, and S1P infusion dose-dependently increased vascular resistance in isolated perfused kidneys from wild-type mice but not mice with compound deficiency in S1PR2&3. Epistatic analysis supported a critical role for S1PR3 in S1P-dependent blood pressure regulation and pointed to a distinct origin of the cardiac phenotype. Although circulating S1P is elevated in hypertensive mice and humans, increasing circulating S1P was not sufficient to induce hypertension in naive mice.

These observations suggests that albumin-S1P crosses the endothelium in resistance arteries to gain access to contractile VSMC S1P receptors, and that myocyte S1PR signaling is essential for vascular resistance and blood pressure maintenance in mice. They also highlight the role for plasma chaperones in specifying vascular responses to S1P and the relevance of S1P as a biomarker and potential therapeutic target for blood pressure regulation and heart failure.

## Introduction

Vascular resistance generated by small arteries and arterioles in peripheral tissues is essential for blood pressure (BP) maintenance and for the distribution of cardiac output (CO). Opening and closing of resistance arteries prioritizes blood supply to organs according to metabolic demand. During physical exercise for example, local vasodilation will increase blood supply to skeletal muscle and myocardium, while sustained or increased constriction in kidneys, liver and gut will decrease supply to those organs, thus limiting demand on CO through its redistribution. Impaired dynamic local control of peripheral resistance is a hallmark of cardiovascular disease and can lead to hypoperfusion, and in severe cases, hypoxia and loss of organ function (1). When systemically increased, peripheral resistance can contribute to hypertension, which is a major risk factor for stroke, myocardial infarction and dementia, represents one of the most important drivers of death and morbidity world-wide, and is not adequately treated with current therapy (2). A systemic decrease in peripheral resistance can induce hypotension, an underappreciated clinical problem with limited treatment options, leading to dizziness and falls and their traumatic consequences, and potentially life-threatening when occurring acutely in the context of systemic infections, trauma, surgery, and allergic reactions, with aggressive treatments forfeiting peripheral perfusion.

A better appreciation of the mechanisms that govern microvascular function may help refine strategies for therapeutic modulation of peripheral resistance and BP. G protein-coupled receptor (GPCR) signaling provides targets for both diagnosis and treatment of cardiovascular disease (3). Knocking down heterotrimeric G proteins in endothelial cells (EC) and vascular smooth muscle cells (VSMC) in mice has established a critical role for GPCR signaling for the regulation of arterial tone and vascular reactivity, and for arterial function in the regulation of BP and the development of hypertension with age and exposure to dietary salt (4–7). The GPCRs acting upstream have not been fully documented. Candidates include receptors that mediate cellular responses to the circulating bioactive lipid sphingosine 1-phosphate (S1P). S1P has long been known to induce both dilatation and constriction of resistance arteries of diverse origin through cognate GPCRs on EC (primarily S1PR1) and VSMC (primarily S1PR2 and S1PR3), respectively (8), and was found to mediate endothelial nitric oxide synthase (eNOS) activation and vasodilation induced by high density lipoproteins (HDL)(9). Mice with EC-selective S1PR1 deficiency (S1PR1 ECKO) present reduced circulating NO derivatives and hypertension (10). Furthermore, mesenteric and cerebral arteries from these mice exhibit reduced flow-mediated dilation (10, 11). Mice deficient in apolipoprotein M (ApoM) and thus in S1P binding to HDL (12) were also hypertensive (13). These experiments confirmed a critical role for S1P in regulating endothelial function and a non-redundant role for HDL-S1P in the activation of EC S1PRs.

The roles of VSMC and cardiomyocyte S1PRs and the sources of S1P activating these receptors are not as well defined. Levels of S1P and its chaperone ApoM are inversely correlated with contractile function and outcome in patients with heart failure (HF) (14, 15), potentially implicating circulating HDL-S1P in the maintenance of cardiac function through actions on endothelial and/or cardiomyocyte receptors. In apparent contrast to the role of S1P in supporting endothelial function, circulating S1P levels are elevated in experimental and clinical hypertension (16, 17) and reduced in patients with trauma and septic shock and in mice with anaphylactic shock (18, 19). This could hint at additional actions of S1P or compensatory regulation of its bio-availability. In accordance with the former, S1P production and S1PR3 signaling were recently implicated in hypertension induced by the glycoprotein sortilin (17). There is also *in vivo* evidence suggestive of a role of S1P in maintaining arterial tone. Although mice deficient in S1PR2 or S1PR3 present with grossly normal BP (10, 20), S1PR2 deficient mice have reduced basal renal and superior mesenteric artery resistance, and arteries from these mice impaired contractile function (20). Increased myogenic tone of resistance arteries observed in heart failure and diabetes has been attributed to induced vessel wall S1P production and S1PR2 signaling (21–23), and global deficiency in S1P production is reported to provide protection against both angiotensin II (AngII)-induced systemic hypertension and hypoxia-induced pulmonary hypertension (24–26). Thus, in addition to promoting vasodilation through S1PR1, S1P signaling has been implicated in cardiac function and vascular resistance. Whether the engagement of myocyte receptors occurs mainly as a result of induced S1P production in the vessel wall or also involves physiological or pathological activation by circulating S1P is unclear, especially considering the barrier imposed by the vascular endothelium.

S1P is synthetized from sphingosine, a metabolite of ceramide, by sphingosine kinases (Sphks) 1 & 2 (encoded by *Sphk1&2*), and can be exported to blood by dedicated transporters (Spns2 and Mfsd2b) (27). S1P is highly abundant in plasma, where it circulates bound to HDL (∼60%), albumin (∼30%) and other minor carriers, also referred to as chaperones (27). The bulk of plasma S1P is provided by erythrocytes and potentially other hematopoietic cells, with possible contribution by ECs (27–29). Platelets also carry large amounts of S1P that can be released upon activation and that distribute mainly to albumin (30), but they do not contribute measurably to plasma S1P under homeostasis (19, 29). We recently reported that S1PR1 signaling on arterial EC is maintained primarily by circulating S1P in naïve mice despite the capacity of EC to produce S1P in stress conditions (28). The dependence of EC S1PR1 on circulating S1P argues against significant production of S1P by vascular cells in naïve mice, although this may change in hypertension and other disease settings (25, 28). Both HDL and albumin can potentially cross the arterial endothelium by paracellular and/or transcellular routes (31), and albumin-S1P can both induce vasoconstriction when administered luminally to resistance arteries and to Langendorff-perfused hearts (32, 33) and improve vascular resistance in anaphylactic shock (19, 34). It is possible that VSMC access is facilitated by compromised barrier function in these settings, but these observations nevertheless suggest the possibility that circulating S1P regulates vascular tone through activation of VSMC receptors also in a basal state.

Circulating S1P levels are high in hypertension and correlate negatively with septic shock severity and heart failure outcomes (14–17). Although causality has been suggested, the consequences of alterations in circulating S1P indiscriminate of its chaperones on BP and cardiac function are not well appreciated, especially in a basal state. In this study we address the consequences of genetically modulating circulating and vessel wall S1P production or the expression of vascular and cardiac S1PRs on vascular resistance, BP, and cardiac function. We observe that while systolic BP increases with age in controls, mice that lack circulating S1P (plasma S1Pless mice) remain normotensive up to 18 months of age and present with compensated left ventricular dysfunction. Bone marrow transplantation and erythrocyte transfusion experiments point to an important role for erythrocyte-derived plasma S1P in the support BP and cardiac function. Analysis of receptor dependence suggested a dominant role for S1PR3 in S1P-dependent regulation of vascular tone and BP and a distinct mechanistic basis for the cardiac phenotype.

## Methods

### Study approval

The study was approved by the animal care and use committee of the Paris Descartes University and by the French Department of Education (02822.02; 03474.02; 03560.02; 28539-202006271229617) and conform to the guidelines from Directive 2010/63/EU of the European Parliament on the protection of animals used for scientific purposes.

### Mice

Mice were housed under specific-pathogen-free conditions in a12-h light and 12-h dark cycle with ad libitum access to water and food (A03-10, Scientific Animal Food and Engineering, Augy, France). Littermate controls of both sexes were used. To assess the effect of circulating S1P deficiency on BP and cardiac function we generated mice lacking Sphk1 and Sphk2 by deletion of *Sphk1&2* in hematopoietic cells with Mx1Cre (induced with 0.5 mg PolyIC twice between p1 and p5). We also generated and characterized mice with *Sphk1&2* deletion in endothelial cells (and a subset of megakaryocytes) with PdgfbiCreERT2 (PdgfbCre), perivascular macrophages (and lymphatic endothelial cells) with Lyve1Cre, myeloid cells with LysMCre, platelets/megakaryocytes (and perivascular macrophages) with Pf4Cre and in VSMC (and cardiomyocytes) with Sm22Cre. Tissue-specific deletion of *S1pr1* in endothelial cells (PdgfbCre) or cardiomyocytes (Myh6Cre) and global *S1pr3*, global or tissue specific (Sm22Cre) *S1pr2* or double *S1pr2/3* knockout mice were used to assess receptor dependency. Conditional knockout alleles for *Sphk1* and *Sphk2* were intercrossed with global knockout alleles in order to increase the efficiency of full gene deletion as previously reported (19, 29, 35).

### Blood and plasma collection

Blood was collected from the retro-orbital venous plexus with heparin- or EDTA-coated capillaries under isoflurane anesthesia. Complete blood cell (CBC) counts were performed with a Hemavet 950 cell counter (Drew Scientific). Blood was centrifugated at 500 g for 10 minutes at room temperature (RT) twice to obtain plasma. Plasma and serum were stored at -80°C for further analysis.

### Sphingolipid measurements by LC-MS/MS

S1P, sphingosine and ceramide quantification in plasma and heart homogenates was performed using a UPLC-MS/MS system consisting of a Waters Acquity and a Triple QuaDripole tandem mass spectrometer or a UHPLC1290 as described previously (36, 37). Individual sphingolipid levels were quantified and normalized to internal standards. Total S1P and ceramide content in the plasma were expressed in μM, while S1P, sphingosine and ceramide in the heart tissue were expressed as pmol/mg of protein.

### Quantification of plasma Renin Angiotensin Aldosterone (RAAS) system components

The RAAS fingerprint was quantified in heparinized plasma by Attoquant Diagnostics (Vienna, Austria) using liquid chromatography-mass spectrometry/mass spectroscopy (LC-MS/MS). Mouse plasma was conditioned for equilibrium analysis at 37 °C followed by enzymatic stabilization as previously described (38). Stabilized equilibrated plasma samples were further spiked with stable isotope-labeled internal standards for both classical and alternative renin-angiotensin system metabolites (angiotensin-(1–10) (Ang I), angiotensin-(1–8) (Ang II), angiotensin-(2–8) (Ang III), and angiotensin-(3–8) (Ang IV)) as well as with internal standards for both steroids, aldosterone-D4 and corticosterone-D4, at a concentration of 200 pg/ml. Internal standards were used to correct for peptide and steroid recovery of the sample preparation procedure for each angiotensin metabolite and steroid in each individual sample. Metabolite concentrations were calculated based on calibration curves in original sample matrix. Ratios of different metabolites were used to calculate angiotensin-converting enzyme (ACE) activity (Ang II/Ang I), plasma renin activity (PRA, Ang I + Ang II), and the adrenal response to Ang II (AA2-Ratio).

### Urine aldosterone quantification

Aldosterone levels in urine were measured at the physiology platform of the Georges Pompidou European Hospital, using liquid chromatography and tandem mass spectrometry (LC-MS/MS) detection to specifically recover urine-free aldosterone after 18-hour acid hydrolysis as previously described (39).

### Biochemical evaluation of kidney function

Proteinuria, blood urea nitrogen (BUN) and serum creatinine were quantified by the UC Davis Comparative Pathology Laboratory.

### Bone marrow transplantation

Wild-type recipient mice were irradiated (6.5 Gy) twice with a 4-hour interval. After 24h, each mouse received 10×10^6^ bone marrow cells (BMC) from sex-matched Sphk1^f/-^: Sphk2^f/-^: Mx1Cre^+/-^ littermate donors (either mixed or in isolation) retro-orbitally under isoflurane anesthesia. Chimeric mice were used 16 weeks post irradiation.

### Red blood cell (RBC) transfusion

Blood was drawn with EDTA-coated capillaries under isoflurane anesthesia into 25% acid citrate dextrose (Sigma) and centrifuged for 15 minutes at 80 g. The platelet-rich plasma and leukocyte-rich buffy coat were removed. Adsol buffer (2g glucose monohydrate, 750g mannitol, 27mg adenine, 90mg NaCl in 100ml final volume), was added to the packed red blood cells at a final concentration of 25%. Packed RBCs were then washed with 15 mL PBS (calcium/magnesium free) and centrifuged for 5 minutes at 100 g. The supernatant was removed, and Adsol solution added to a final volume 25%. Cells from different donors were then pooled and maintained at a density of 4×10×9 to 8×10×9 cells/mL. Recipient mice received 350 uL of packed RBCs by intravenous (i.v.) injection. Cardiac function was assessed before and 48 hours post-transfusion and BP before and 3 consecutive days after transfusion in separate cohorts of mice.

### Losartan administration

Losartan (20mg/kg) was administered by intraperitoneal (i.p.) injection for 3 consecutive days. BP was recorded by telemetry for 24h before the first injection and 24h after the third injection.

### Angiotensin II infusion by osmotic mini-pumps

Hypertension was induced by infusion of AngII (1500 ng/kg/min for 7 days of 500 ng/kg/min for 28 days, as indicated) with osmotic mini-pumps (Alzet Model 2004) implanted subcutaneously (s.c.) in naïve mice. BP was recorded by telemetry before and either continuously for 7 consecutive days (high dose AngII) or for 24h on day 0, 7, 14, 21 and 27 after implantation of minipumps (low dose AngII).

## L-Name administration

L-Name (Tocris) was administered in the drinking water (1 g/L) for 7 consecutive days. BP was recorded by telemetry for 24h from day 0 to 7 after L-NAME administration.

### BP measurements

#### Acute carotid artery BP measurements

Mice were injected i.p. with buprenophrine (0.1mg/kg in NaCl 0.9%) 20 min before surgery. They were subsequentially anesthetized with isoflurane (4 % induction; 2% stabilization) and catheter connected to a pressure transducer implanted on the left carotid artery to acquire BP and heart rate (HR). Diastolic/systolic arterial pressure and heart rate were recorded with Biopac student lab software under 2% isoflurane. Basal or post treatment BP was monitored acutely for 20 minutes at baseline or after treatment with values reported every 60 seconds. BP is expressed as average during 15-20 minutes of stable recording.

#### Telemetry based aortic BP recordings

Systolic BP (SBP), diastolic BP (DBP) and HR were measured in conscious mice using Data Sciences International (DSI) implantable radiotelemetry transmitters. The analgesic (as described above) was administered i.p. 20 minutes before surgery and s.c. 6 hours after surgery. Mice anesthetized with ketamine /xylazine (100/10 mg/kg) were implanted with carotid artery catheters through the aortic arch and radio transmitters (PA-C10) implanted in a dorsal s.c. pocket. After 10-12 days of recovery, BP was monitored continuously with values reported every 60 seconds. Basal BP was measured for at least 24 hours for up to 3 consecutive days. BP recordings are expressed as the average of 10-11 hour intervals (8 pm-7 am; 8 am-6 pm). Ponemah 6.x software was used to record and analyze BP.

#### Tail cuff plethysmography

Non-invasive SBP measurements were performed by tail-cuff plethysmography (BP-2000-M4 Blood Pressure Analysis System™, Visitech Systems). Mice were acclimatized and then placed in a preheated chamber with a pulse sensor around their tails for BP recordings. Three to six consecutive days of training were performed before basal BP was recorded for three consecutive days. The average SBP was calculated from a minimum of 10 measurements for each recording session, and the average 3 days of recording presented.

#### Dietary salt challenge

Mice were fed with normal chow diet (ND; 0,3% Na+, 0,8% K+), low sodium diet (LSD; 0,03% Na+; 0,8% K+) or high sodium diet (HSD; 4% Na+, 0,8% K+) for 3 weeks. They were then housed in metabolic cages with free access to water for 14 hours for urine collection. For each mouse, BP measurements as well as blood and urine collection were made at baseline (ND) and at day 19, 20 and 21 of the diets (HSD/LSD), respectively. BP was measured by tail-cuff plethysmography. Blood levels of sodium (Na), potassium (K), ionized calcium (iCa), glucose (Glu), hematocrit (Hct), hemoglobin (Hb), as well as pH, bicarbonate (HCO3), carbon dioxide partial pressure (PCO2), total carbon dioxide (TCO2), oxygen partial pressure (PO2), base excess (BE) and oxygen saturation (sO2) were measured or calculated using an i-STAT CG8+ cartridge analysis kit within 10 minutes from collection.

#### Kidney isolation, perfusion and vascular resistance measurements

In anesthetized mice, the renal artery was cannulated and the kidney was excised for perfusion with PSS (135 mmol/L NaCl, 15 mmol/L NaHCO3, 4.6 mmol/L KCl, 1.5 mmol/L CaCl2, 1.2 mmol/L MgSO4, 11 mmol/L glucose, 10 mmol/L N-2-hydroxyethylpiperazine-N-2-ethylsulfonic acid (HEPES) at 37°C as described previously (40). Perfusion was maintained at a rate of 600 µL/min and monitored using a PM-4 pressure transducer (LSI, Burlington, VT, USA). Dilation to acetylcholine (1 µmol/L) after pre-contraction to phenylephrine (1 µmol/L) was used to evaluate the integrity of the endothelium. Renal vascular reactivity was tested using KCl (80 mmol/L). The basal perfusion flow rate was then increased step-wise in order to determine the flow-pressure relationship. Concentration-response curves to S1P reconstituted with lipid free BSA (BSA-S1P) were performed.

#### Ultrasound-based assessment of left ventricular systolic function, cardiac output and peripheral hemodynamics

Ultrasound examination was performed under light anesthesia (0.75 % isoflurane) on mice placed on a temperature-controlled heating platform using an echocardiograph (Acuson S3000, Erlangen, Germany) equipped with a 14-MHz linear transducer (14L5 SP) as previously described (40). Pulmonary artery diameters were acquired by a left parasternal long-axis B-mode image and a flow profile recorded by a pulsed Doppler sample on the longitudinal axis. Blood flow velocities were measured and corrected for the angle between the long axis of the artery and the Doppler beam. Wall thickness and internal diameters in systole and diastole were measured in M-mode. Left ventricle (LV) end-systolic (ESV) and end-diastolic (EDV) volumes, as well as ejection fraction (EF) were calculated according to Teichholz ((7.0 / (2.4 + LVIDd)) * LVIDd×3 * 100%), and the LV mass was calculated according to Troy (1.05 ([LVIDd + PWTd + IVSTd]3-[LVIDd]3) g). Cardiac output (mL/minute) was calculated from the left ventricular diastolic and systolic volumes (SV (stroke volume) x HR (heart rate)) or from the mean pulmonary blood flow velocities and pulmonary diameter as described (40). Both approaches provided consistent results. All measures were averaged from at least three cardiac cycles. Peak systolic, end-diastolic and time-averaged mean blood flow velocities (mBFVs) of the right renal artery (RRA) and superior mesentery artery (SMA) were measured in color-Doppler mode with angle correction. The operator was blinded to genotype.

#### Cardiac stress testing by dobutamine

Heart rates, peak systolic, end-diastolic parameters were measured at baseline and 10 minutes after intraperitoneal dobutamine administration (1,5 µg/kg) to evaluate cardiac reserve capacity.

#### Immunohistochemistry

To assess cardiac remodeling, hearts were perfused, excised and fixed for 1h in 4 % PFA at RT before paraffin embedding. Hearts were cut into 5μm thick sections. Picrosirius Red staining (the collagen fibers were stained red and myocytes yellow), were performed for myocardial fibrosis evaluation. The degree of fibrosis was calculated as the ratio of the total area of fibrosis to entire area of visual field of the section using ImageJ.

#### Immunofluorescence staining and imaging

Immunofluorescent staining was performed as previously described (28). Briefly, after transcardial perfusion of heparinized DBPS and 1% PFA, tissue (heart, adrenal gland) was post-fixed for 1 hour in 4 % PFA at RT placed in 30% sucrose overnight, and 10 µm cryosections were prepared. Cryosections were blocked for 2h at RT with Blocking buffer. Thereafter, primary antibodies diluted 1:1 in BB and PBS were incubated overnight at 4°C whereas secondary antibodies for 2 hours at room temperature. Frozen sections were mounted in Fluoromount-G (#0100-01, Southern Biotech). For kidney histology, sections were then stained with Hematoxylin and Eosin (H&E) (Bio-optica Fast H&E kit #04-061010) following manufacturer’s instructions. The following primary antibodies were used: mouse anti-ASMA-Cy3 (1:250, #C6198, Sigma), goat anti-CD31 (1:300, #AF3628, R&D Systems), rabbit anti-Lyve1 (1:300, #11034, Angiobio), rat anti-CD206 (1:200, #MCA2235GA, Bio-Rad), rabbit anti-Cardiac Troponin T-Alexa647 (1:100 #EPR20266) Corresponding secondary antibodies were purchased from ThermoFisher (donkey anti-primary antibody coupled with AlexaFluor488, AlexaFluor555 or AlexaFluor647 and streptavidin conjugated with AlexaFluor405) or Jackson ImmunoResearch (donkey anti chicken AlexaFluor488 and donkey anti-rabbit AlexaFluor405). Confocal images were acquired on an inverted Leica SP8 laser scanning microscope using LAS-X software (Leica, Germany) with 20x (NA 0.75) or 40x (NA 1.30) oil immersion objectives. Single and Z-stack images were processed equally with LAS-X software and ImageJ/Fiji (National Institutes of Health, USA). Analyses were performed with CellProfiler (version 3.9.0 – 4.2.1, Broad Institute, Boston, MA, USA)

#### Immunofluorescence imaging and quantification of adrenal glands

10 µm adrenal cryosections were permeabilized with TBS 0.1% Triton X100 for 15 minutes. Then sections were then blocked with 5% BSA and 10% normal donkey serum (NDS) diluted in TBS 0.05% Triton X100 for 2 hours at room temperature. Rabbit anti-Cyp11b2 (mCyp11b2, 1/100, generous gift from CE Gomez-Sanchez) was diluted in 3% BSA and 3% NDS in TBS 0.05% Triton X100 and incubated overnight at 4°C. Cyp11b2 was detected with donkey anti-rabbit Alexa 647 in 3% BSA and 3% NDS in TBS 0.05% Triton X100. Nuclei were counterstained using 4′,6-diamidino-2-phenylindole (DAPI) (1/5000, Roche Diagnostics GmbH). Whole slide imaging was performed using the Olympus Slideview VS200 digital slide scanner (Olympus Corporation, Tokyo, Japan) with UPlanXApo 20x/0.80 objective lens. Image analysis was performed using HALO software (version 3.6.4134, Indica Labs) with the HighPlex Fl v4.2.14 module. Background correction was applied to reduce non-specific fluorescence and autofluorescence in the tissue. ROIs corresponding to the adrenal tissue were manually annotated. The adipose tissue around the adrenal and the medulla was manually removed and excluded from the analysis. DAPI was used for traditional nuclear segmentation within the ROIs. Cyp11b2 positive cells were identified using Alexa Fluor 647.

#### Wheat germ agglutinin (WGA) staining

Mouse hearts were fixed with 4% paraformaldehyde overnight at 4°C, cut in two parts and embedded in paraffin for sectioning. Myocardial sections of 5µm were stained with 40 μg/mL WGA (W7024; Invitrogen) in PBS for 2 hours at room temperature in order to label cardiomyocyte membranes. Nuclei were counterstained with Hoechst. Immunofluorescence images of heart sections were captured with an inverted Leica SP8 laser scanning microscope using LAS-X software (Leica, Germany). Cross-sectional area was analyzed with ImageJ.

#### Fluorescent albumin leakage

Vascular permeability was assessed by i.v. injection of fluorescence labelled albumin (BSA-AlexaFluor647, 20mg/kg, #A34785, ThermoFisher) under isoflurane (3%) anaesthesia. After 1h, mice were transcardially perfused with heparinized DPBS followed by 1% PFA and mesenteric arteries were collected and post-fixed in 4% PFA overnight at 4°C and washed three times in PBS. The tissue was prepared for cryosections as described above.

#### Real-Time Quantitative PCR (RT-qPCR)

Mouse total RNA purification was performed using the miRNeasy Mini Kit (Qiagen, Hilden, Germany). RNA yield was measured using the NanoDrop 2000 (Thermo Fisher Scientific, Wilmington, USA). Quantitative RT-PCR was performed with cDNA at a concentration of 100 ng per 10 µl reaction volume. All quantitative PCR experiments were performed using SyberGreen Gene Expression Assays (please see Table S2 for details) with a CFX96 Real-Time PCR Detection System (BioRad, Hercules, California, USA). The 2×-(ddct)-method was used for fold change calculation.

#### RNAseq profiling and analysis

Total RNA was isolated from cardiac tissue using TRIzol Reagent and RNeasy mini kits (QIAGEN) following the manufacturer’s protocol. RNA quality was assessed with Nanodrop. Library preparation and sequencing were performed Biomics Platform, C2RT, Institut Pasteur, (Paris, France). Quality control was performed on a Fragment Analyzer (Agilent). Sequencing was performed on a NextSeq2000 (Illumina) to obtain 68 base single-end reads. The RNA-seq analysis was performed with Sequana 0.14.1 using the RNA-seq pipeline 0.15.2 (https://github.com/sequana/sequana_rnaseq) built on top of Snakemake 7.8.5 (41). Briefly, reads were trimmed from adapters using Fastp 0.20.1 (42) then mapped to the Mus musculus GRCm39 genome assembly from Ensembl using STAR 2.7.8a (43). FeatureCounts 2.0.1 (44) was used to produce the count matrix, assigning reads to features using corresponding annotation v109 from Ensembl with strand-specificity information. Quality control statistics were summarized using MultiQC 1.11 (45). Statistical analysis on the count matrix was performed to identify differentially regulated genes. Clustering of transcriptomic profiles were assessed using a Principal Component Analysis (PCA). Differential expression testing was conducted using DESeq2 library 1.30.0 (46) scripts indicating the significance (Benjamini-Hochberg adjusted p-values, false discovery rate FDR < 0.05) and the effect size (fold-change) for each comparison. Finally, enrichment analysis was performed using modules from Sequana. The GO (Gene Ontology Consortium; (47)) enrichment module uses PantherDB (48) and QuickGO (49) services. The KEGG pathways enrichment uses GSEApy 0.10.8 (50), EnrichR (51) and KEGG database (52). Programmatic accesses to online web services were performed via BioServices 1.10.1 (53). Only significant enrichments with a Benjamini Hochberg-adjusted P value of less than 0.05 were considered.

#### Statistical analyses

Statistical analyses were performed with Prism 10 software (Prism, GraphPad, San Diego, USA). A value of *P*<0.05 was considered statistically significant. Specific statistical tests performed are indicated in Table I. Parametric tests were applied only if normal distribution could be confirmed. Each symbol in graphical representations show data or pooled quantification from one mouse. Scatter dot plot show mean ± SEM.

## Results

### S1P maintains BP and contributes to its increase with age in mice

In accordance with an essential role for S1PR1 in maintaining endothelial function *ex vivo* (10, 11), selective loss of either HDL-associated S1P or EC S1PR1 render mice hypertensive (10, 13). On the other hand, normal BP in mice lacking S1PR2 or S1PR3 (10, 34) may suggest that support of myogenic tone by S1P or its receptors observed *ex vivo* (10, 54, 55) is redundant in the physiological context. To challenge this notion, we measured BP in mice with loss of both HDL- and albumin-associated S1P generated by postnatal deletion of *Sphk1&2* with an Mx1 promoter-driven inducible Cre recombinase *(Sphk1^f/f^:2^-/-^:*Mx1Cre^+^) (29). Plasma S1P levels in these mice (AKA as plasma S1Pless; (35)) range from undetectable to ∼20 nM, i.e. around 2 % of littermate controls (19, 28, 29, 37). In direct contrast to mice with selective deficiency in HDL-S1P, systolic (S), diastolic (D) and mean (M) BPs in male plasma S1Pless mice were ∼15-20 mmHg lower than littermate controls as assessed under anesthesia with aortic BP probes directly connected to pressure monitors (Fig. 1A). A similar reduction in BPs was observed in plasma S1Pless mice generated through a breeding strategy that used conditional knockout alleles for both Sphks (*Sphk1^f/-^:2^f/-^:*Mx1Cre^+/-^) (Fig. 1B) (19, 28). Female plasma S1Pless mice generated with this strategy were also hypotensive (Fig 1C). Lower BPs in plasma S1Pless mice was also observed by non-invasive tail-cuff plethysmography in restrained unsedated males (Fig 1D) and by twenty-four-hour telemetry recordings during light and dark cycles in freely moving males (Fig 1E). Telemetry recordings also revealed a blunted day-night BP increase (Fig 1E). No differences were observed in HR by any method in either sex (Fig 1A-C,E). The impact of S1P on BP was age dependent: while SBP of control mice increased with age, this was not the case in plasma S1P deficient mice (Fig 1F; Supplemental Fig 1). Thus, S1P maintains BP in naïve male and female mice on standard diet and contributes to the increase in BP with age.

**Figure 1.**
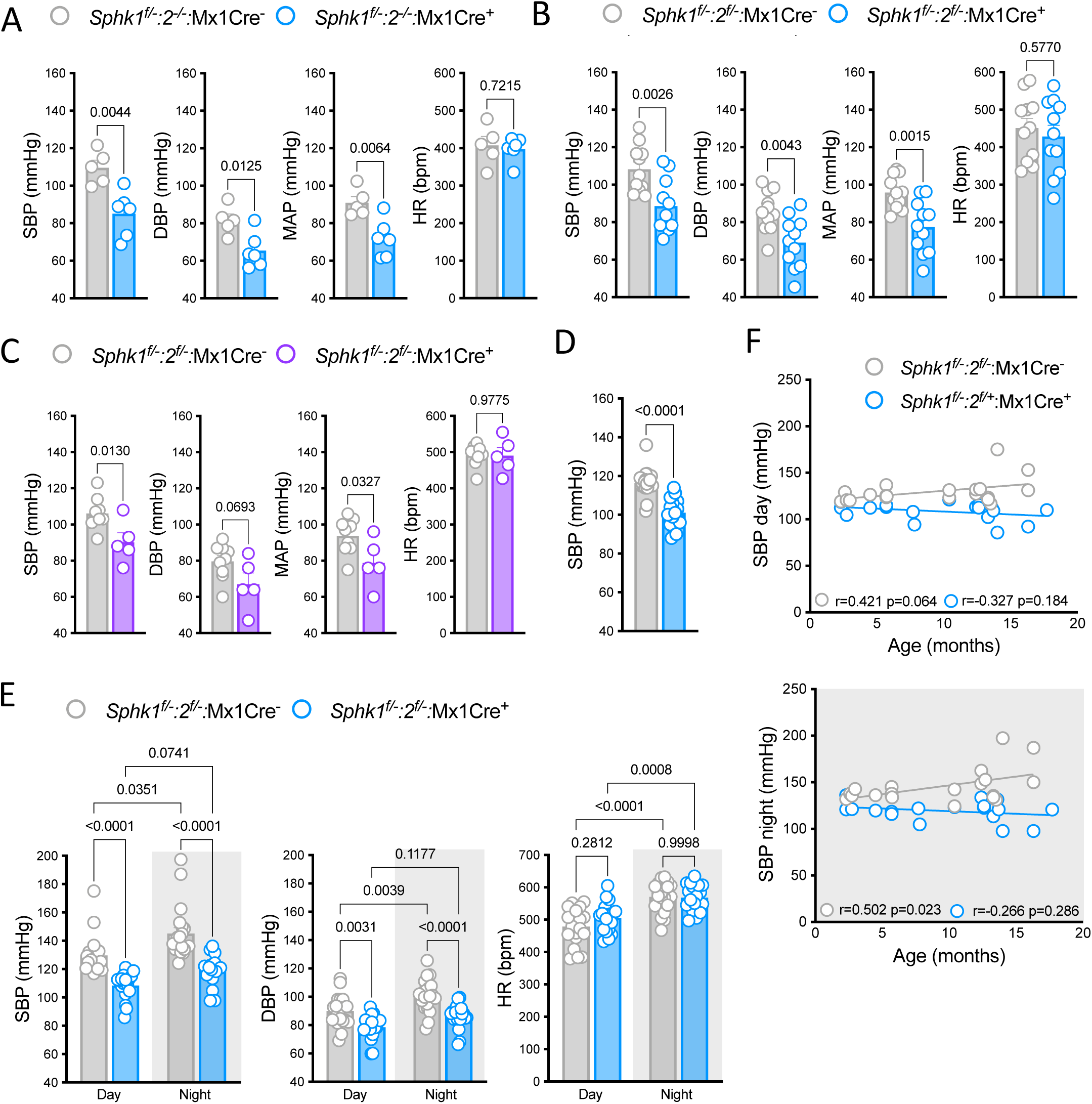
Plasma S1Pless mice have lower BP than age and sex-matched littermate controls. (**A-C**). Invasive central BP and HR measurements of anesthetized *Sphk1 ^f/f^: 2^-/-^:* Mx1Cre^+^ (A) (n=6) and *Sphk1 ^f/-:^2 ^f/-^:* Mx1Cre^+^ (B) (n=11) males and *Sphk1 ^f/-:^2 ^f/-^:* Mx1Cre^+^ female (C) (n=5) plasma S1Pless mice and their respective age and sex-matched Mx1Cre^-^ littermates (n=5,11,9). Non-invasive tail cuff plethysmography-based SBP measurements of *Sphk1 ^f/-:^2 ^f/-^:* Mx1Cre^+^ (n=18) plasma S1Pless and Mx1Cre^-^ control males (n=20). (**E**) Dark and light cycles invasive radiotelemetry-based central BP and HR measurements of *Sphk1 ^f/f^:2^-/-^:* Mx1Cre+ (n=18) plasma S1Pless and littermate control males (n=18). (**F**) Correlation between age and SBP in plasma S1Pless mice and littermate control mice presented in D (upper panel) and E (lower panel, nocturnal SBP only shown). Each dot represents one mouse. Scatter dot plots show mean±SEM. Statistical analysis was performed using unpaired t-test or Mann-Whitney; two-way ANOVA or Kruskal-Wallis followed by Dunnett’s or Sidak’s multiple comparison and Pearson correlation.

### Erythrocyte-derived circulating S1P maintains BP in mice

The Mx1Cre driver used to generate plasma S1Pless mice provides efficient inducible excision in the hematopoietic compartment, which provides the bulk of plasma S1P (28, 29), but also targets vascular and other cell types in which Sphk activity has been implicated in myogenic tone and BP regulation (23, 28, 35, 54, 56, 57). Moreover, loss of S1P production in lymphatic endothelial cells (LEC) in this model impairs lymphocyte trafficking and is predicted to impact lymphatic vascular development (28, 29, 58), both with potential impact on BP regulation (59, 60). To dissect the Mx1Cre-sensitive sources of S1P implicated in the observed BP phenotype, we measured BP in >5 month old anesthetized male mice with selective deficiency in S1P production in EC (*Sphk1*^f/-^:*2*^f/-^:PdgfbCreERT2, perinatal induction), VSMC and cardiomyocytes *(Sphk1*^f/-^:*2*^f/-^:Sm22Cre) or LEC *(Sphk1*^f/-^:*2*^f/-^:Lyve1Cre) with Cre drivers validated for excision efficiency ((28); Supplemental Fig 2). Disabling S1P production in the cells targeted by these Cre drivers did not reduce BP (Fig 2A-C). To specifically address the role for hematopoietic S1P sources, we replaced Sphk deficient bone marrow cells (BMC) of lethally irradiated young S1Pless adults with wild-type BMC. If anything, BPs in these mice were higher than littermate controls (Fig 2D). Like plasma S1P production (28, 29), the BP phenotype therefore tracked with the hematopoietic compartment. While erythrocytes provide the bulk of plasma S1P, platelets and monocytes/macrophages may also release S1P in certain contexts (61). Yet selective impairment of S1P production in platelets *(Sphk1*^f/f^:*2*^-/-^: Pf4Cre)(19, 62) or myeloid cells *(Sphk1*^f/-^:*2*^f/-^: LysMCre) (28)) did not impact BP (Fig 2E,F), pointing to a role for erythrocyte-derived plasma S1P. Substantiating the above observations, irradiated wild-type recipients of S1P deficient BMC (*Sphk1*^f/-^:*2*^f/-^:Mx1Cre^+^) had lower BP than recipients of BMC from controls (*Sphk1*^f/-^:*2*^f/-^:Mx1Cre^-^) (Fig 2G), and transfusion of erythrocytes from wild-type donors (Fig 2H) eliminated the difference in SBP between plasma S1Pless mice and littermate controls (Fig 2I). S1P levels in plasma isolated 5 days post-transfusion were 5-fold higher than levels pre-transfusion, consistent with a partial normalization of plasma S1P levels (Fig 2J). These observations indicate that S1P production by hematopoietic cells - primarily erythrocytes - is essential for BP maintenance in naïve mice.

**Figure 2.**
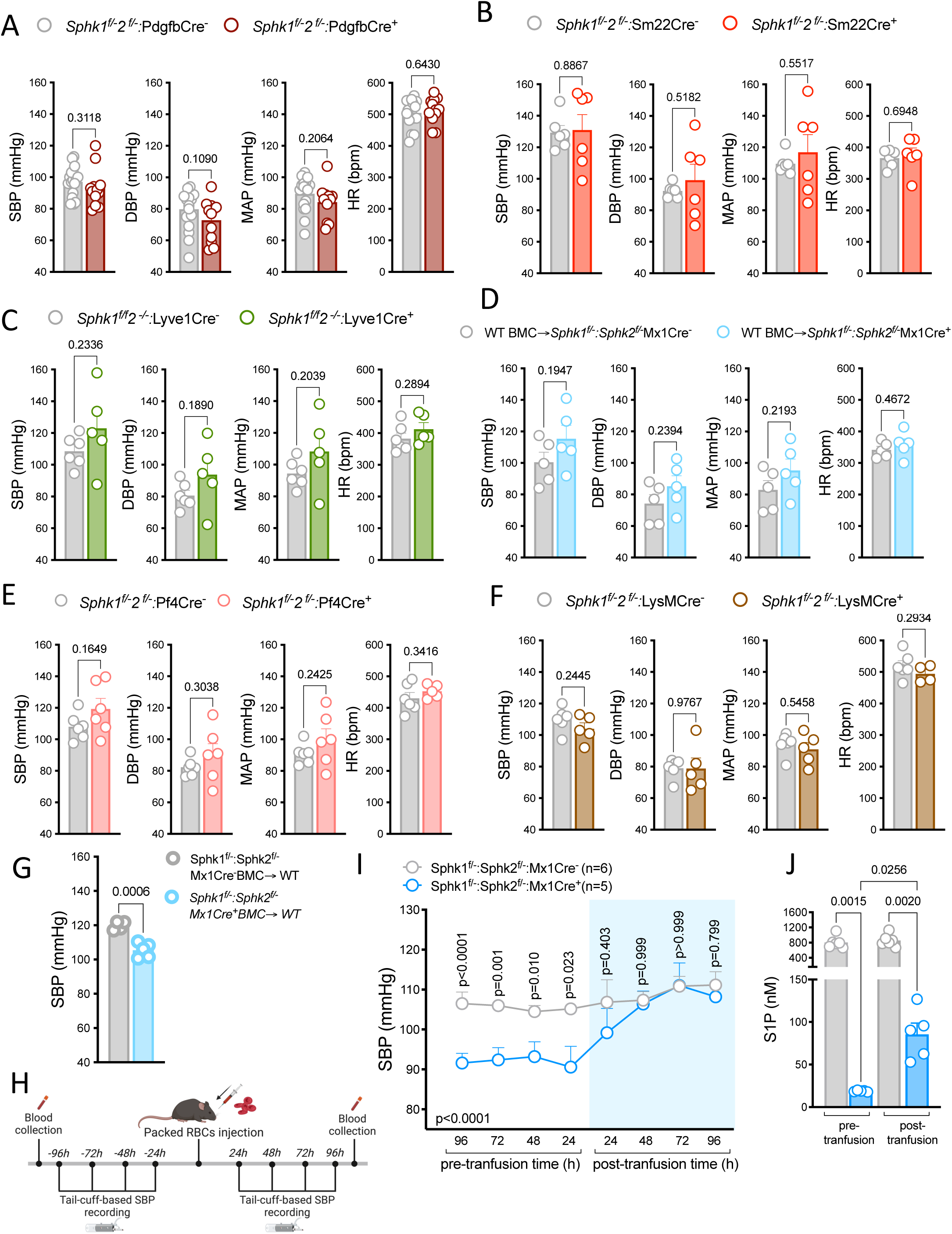
Erythrocyte-derived S1P maintains BP. (**A-F**) Invasive central BP and HR measurements of anesthetized male mice lacking S1P production in endothelial cells (*Sphk1^f/-^:2^f/-^:* PdgfbCre^+^; n=12; A), VSMC and cardiomyocytes (*Sphk1^f/-^:2^f/-^:*Sm22Cre^+^; n=6; B), lymphatic endothelial cells and perivascular macrophages (*Sphk1 ^f/f^:2^-/-^:* Lyve1Cre^+^;n=5; C), non-hematopoietic cell targets of Mx1Cre recombinase (*Sphk1*^f/-^:*2*^f/-^:Mx1Cre^+^ mice transplanted with wild type bone marrow cells (BMC); n=5; D), platelets and perivascular macrophages (*Sphk1^f/-^:2^f/-^:*Pf4Cre^+^; n=6; (F), myeloid cells (*Sphk1^f/-^:2^f/-^:*LysMCre^+^; n=5; E), and their respective littermate controls (n=6) (**G**) Tail-cuff based SBP measurements of wild-type mice transplanted with BMC capable (*Sphk1*^f/-^:*Sphk2*^f/-^:Mx1Cre^-^ n=4) or not (*Sphk1^f/-^:2^f/-^*:Mx1Cre^+^n=5) of S1P production. (**H**) Experimental design and timeline of erythrocyte transfusion and BP measurements in I and J. (**I**) Tail-cuff based SBP in plasma S1Pless (*Sphk1*^f/-^:*2*^f/-^:Mx1Cre^+^; n=5) and control (*Sphk1*^f/-^:*Sphk2*^f/-^:Mx1Cre^-^; n=6) males pre and post erythrocyte transfusion. (**J**) Analysis of plasma S1P levels in mice shown in I two weeks before and 5 days post erythrocyte transfusion. Each symbol represents one mouse in bar graphs; Scatter dot plots show mean±SEM. Statistical analysis was performed using unpaired t-tests (A-G), one-way (J) or two-way (I) ANOVA followed by Sidak’s multiple comparison.

### S1P maintains BP by supporting peripheral vascular resistance

S1P induces both endothelium-dependent dilation and VSMC-mediated constriction of isolated arterial segments examined *ex vivo* (8, 10). Although dilatory effects dominate when physiological concentrations of albumin-S1P are infused into the lumen of pressurized uterine and mesenteric resistance arteries, constriction is observed at higher doses or when eNOS is impaired (32). In Langendorff-perfused murine hearts, infusion of albumin-S1P substantially reduces coronary blood flow at physiological concentrations (63). Thus, albumin-S1P can cross the endothelium for access to VSMC receptors in vessel/organ preparations *ex vivo*. Suggesting access also *in vivo,* we observed an accumulation of Alexa647 both within the endothelium and in Lyve1+ macrophages outside the VSMC layer of mesenteric arteries 60 minutes after intravenous administration of Alexa647-labelled albumin to naïve wild-type mice (Supplemental Fig 3A). This contrasts lack of detectable extravasation in cerebral arterioles (28), and suggests the possibility that plasma S1P maintains vascular tone by continuous activation of VSMC S1PRs. Supporting this notion, Echo Doppler analysis revealed elevated right renal and superior mesentery artery blood flow velocities that coincided with a reduction in total peripheral resistance in plasma S1Pless mice relative to littermate controls (Fig 3A). CO (M-mode derived in Fig3B; velocity derived in Supplemental Fig3B) did not differ between the genotypes. Intriguingly, rather than a decrease in renin– angiotensin–aldosterone system (RAAS) activity, plasma S1Pless males displayed a substantial increase in equilibrium angiotensin peptides (Fig 3C). Plasma renin activity was calculated to be higher, and angiotensin-converting enzyme activity slightly elevated (Fig 3D). Renal architecture and renin expression were not impacted by S1P deficiency, nor were blood urea nitrogen/serum creatinine ratios or creatine kinase levels (Supplemental Fig 3C-F). Plasma S1Pless mice also did not display proteinuria even at >1 year of age (Supplemental Fig 3F). Plasma corticosterone and aldosterone levels also did not differ between genotypes (Fig 3F), and no change was seen in the abundance of aldosterone synthase positive cells within the adrenal cortex (Supplemental Fig 3G-H). Grossly normal renal and adrenal structure and function and higher RAAS activity may suggest that the RAAS is engaged to compensate for loss of S1P-mediated support of vascular tone. Accordingly, plasma S1Pless females, which had a less consistent BP reduction than the males, displayed an even greater increase in angiotensin peptides and renin activity (Supplemental Fig 3I). Moreover, when we treated S1Pless and littermate control mice with an angiotensin receptor blocker (Losartan; 20mg/kg; i.p) for 3 consecutive days and monitored changes in BP (64), plasma S1Pless mice showed a greater drop in SBP during both light and dark cycle (Fig 3F). These observations argue that plasma S1P maintains BP by vascular tone regulation, and that the impact of S1P deficiency is partially compensated for by increased RAAS activity.

**Figure 3.**
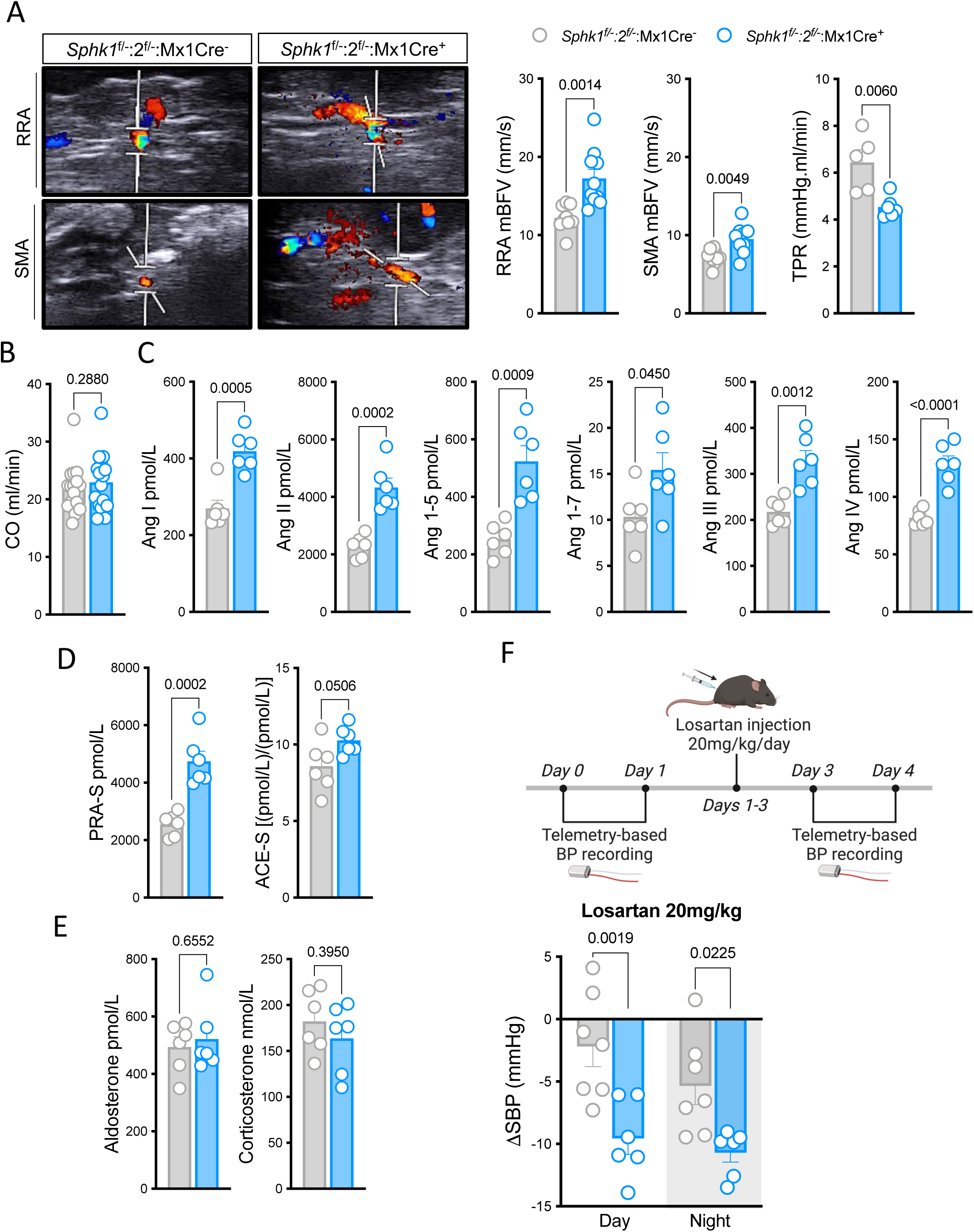
Plasma S1P deficiency result in decreased peripheral vascular resistance. (**A**) Mean blood flow velocities (mBFVs) measured by ultrasound in right renal artery (RRA) and superior mesentery artery (SMA) of plasma S1Pless (*Sphk1*^f/-^:*Sphk2*^f/-^:Mx1Cre^+^; n=10) and littermate control (*Sphk1*^f/-^:*Sphk2*^f/-^:Mx1Cre^-^; n=9) males. Total peripheral resistance (TPR) was calculated as MAP measured by acute pressure response divided by pulmonary artery mBFV (n=5-6). Left panel: Representative doppler images from RRA and SMA. (**B**) Cardiac output (CO) of of plasma S1Pless (*Sphk1*^f/-^:*Sphk2*^f/-^:Mx1Cre^+^; n=16) and littermate control (*Sphk1*^f/-^:*Sphk2*^f/-^:Mx1Cre^-^;n=17) males. (**C**) Equilibrium angiotensin metabolite profile of plasma S1Pless (*Sphk1*^f/-^:*Sphk2*^f/-^:Mx1Cre^+^; n=6) and littermate control (*Sphk1*^f/-^:*Sphk2*^f/-^:Mx1Cre^-^;n=6) males. (**D**) Renin activity (PRA-S), calculated as the sum of AngI+AngII, and ACE activity (ACE-S) as ratio between Ang II and Ang I. (**E**) Mass spectrometry–based determination of aldosterone and corticosterone. **(F**) Telemetry based night and day delta SBP after 3 consecutive days of Losartan (20 mg/kg i.p.) treatment of Plasma S1Pless (*Sphk1*^f/-^:*Sphk2*^f/-^:Mx1Cre^+^; n=6) and littermate control (*Sphk1*^f/-^:*Sphk2*^f/-^:Mx1Cre^-^; n=7) males. Each symbol represents one mouse. Experimental design and timelidne shown in upper panel. Scatter dot plots show mean±SEM. Statistical analysis was performed using unpaired t-test or Mann-Whitney and two-way ANOVA followed by Sidak’s multiple comparison.

### S1P promotes vascular resistance and sustains BP via S1PR2&3

We next addressed the implication of S1PRs in vascular tone and BP regulation by plasma S1P. Considering the evidence for a role for S1PR1 in supporting endothelial function and NO-dependent BP regulation, we first addressed the possibility that Sphk deficiency in hematopoietic cells increased S1PR1-mediated vasodilation by increased abundance of sphingosine as a substrate for vessel wall Sphks. However, supply of L-NAME to the drinking water of plasma S1Pless males for 7 days did note eliminate, but instead exacerbated the difference in BP with littermate controls (2 of which died after 5 days of treatment; Fig 4A). Rather than hyperactivation of endothelial S1PR1 signaling, this pointed to a role for VSMC S1PRs partially masked by a loss of S1PR1-mediated vasodilation predicted with deficiency in plasma S1P (10, 28). As we observed reduced renal vascular resistance in S1Pless mice *in vivo*, we employed a renal perfusion system which requires interaction between myogenic tone and flow-mediated dilation to investigate the role of S1PRs in modulating renal vascular resistance *ex vivo* (Fig 4B). Infusion of albumin-S1P into phenylephrine pre-constricted kidneys from wild-type mice resulted in an initial drop in renal vascular resistance demonstrated by a decrease in perfusion pressure, followed by a dose-dependent increase in vascular resistance starting in the low nanomolar range (Fig 4C). The initial drop in vascular resistance was also observed with S1P free albumin and persisted in mice lacking S1PR1 on endothelial cells (Fig 4C,D). Vasoconstriction induced by increasing doses of albumin-S1P was significantly blunted in kidneys from S1PR3 deficient mice, slightly reduced in kidneys from S1PR2 deficient mice, and fully abolished in kidneys from mice with compound deficiency in the two receptors (Fig 4D). These data suggest that albumin-S1P promotes an S1PR2/3-mediated constrictor response at physiological concentrations with no significant contribution of VSMC S1PR1 (63).

**Figure 4.**
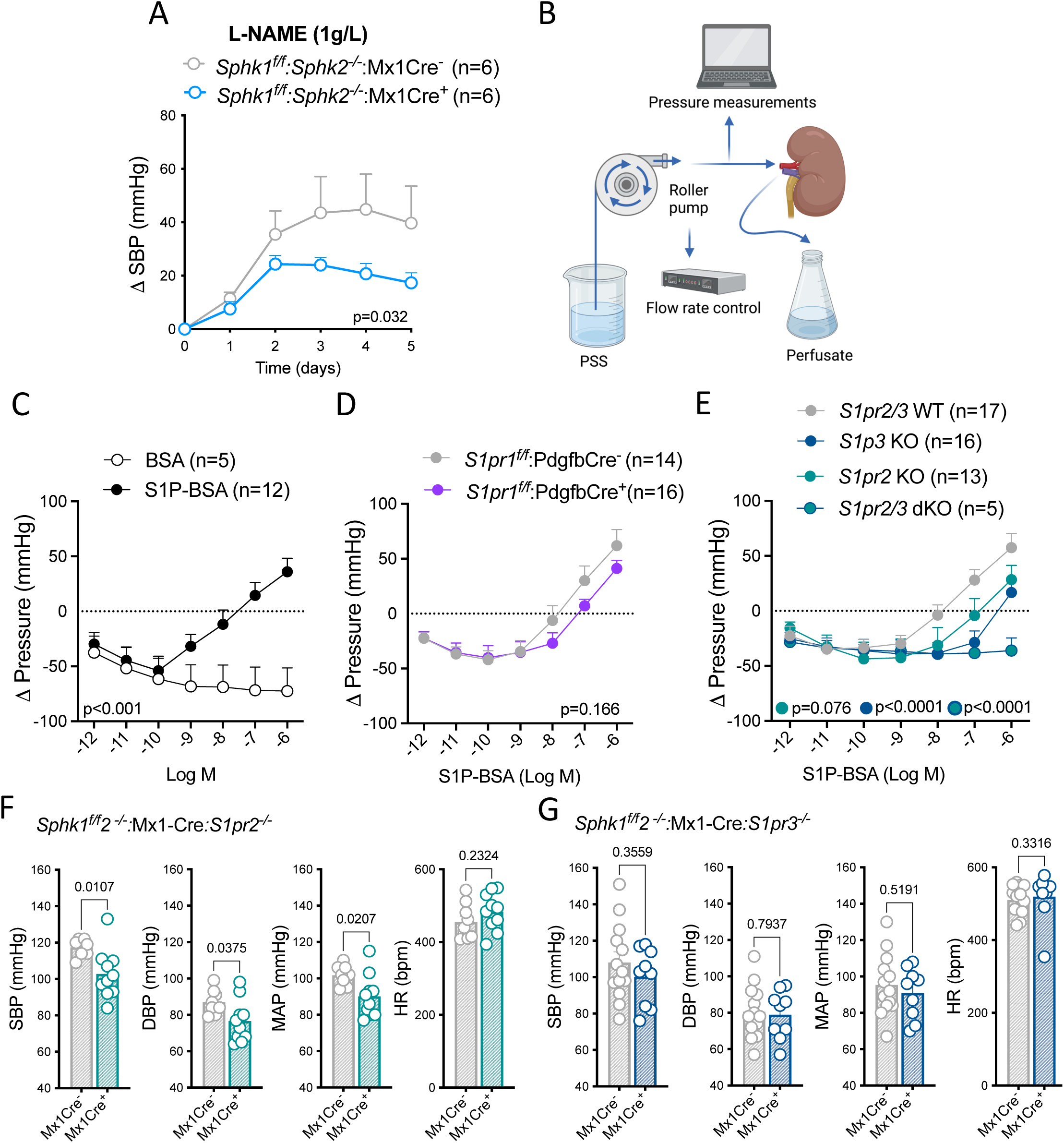
Plasma S1P sustain vascular tone mainly through S1PR3 activation. (**A**) Telemetry based delta SBP before and during 7 days of L-NAME (1g/L) treatment in the drinking water of plasma S1Pless (*Sphk1*^f/f^:*2*^-/-^:Mx1Cre^+^ ; n=6) and littermate controls (*Sphk1*^ff^:*2*^-/-^:Mx1Cre^-^; n=6). Note that as two control mice died on day 6 of treatment only the first 5 days are shown. (**B**) Schematic illustration of the kidney perfusion system used in C-D. **C**. Delta pressure with increasing concentrations of BSA (n=5) or S1P-BSA (n=12) infusion in wild-type kidneys. (**C**) Delta pressure with increasing concentrations of S1P-BSA in S1pr1 ECKO (*S1pr1*^f/f^:PdgfbCre^+^ ; n=16) and littermate control (*S1pr1*^f/f^:PdgfbCre^-^; n=14) kidneys. (**D**) Delta pressure with increasing concentrations of S1P-BSA in kidneys from S1pr2 deficient (KO) (n=13), S1pr3 deficient (KO) (n=16), S1pr2/3 deficient (dKO) (n=5) and shared littermate control (n=18) mice. **(E)** Arterial pressures of anesthetized plasma S1Pless males in an S1PR2 deficient background (*Sphk1^f/f^*:*2^-/-^*: Mx1Cre^+^:*S1pr2^-/-^*; n=9) and of their respective littermate controls (*Sphk1^f/f^:2^-/-^:* Mx1Cre^-^:*S1pr2^-/-^*; n=8). (**F**) Arterial pressure of anesthetized plasma S1Pless males in an S1PR3 deficient background (*Sphk1^f/f^:2^-/-^:* Mx1Cre+:*S1pr3^-/-^*; n=9) and of their respective littermate controls (*Sphk1^f/f^:2^-/-^*: Mx1Cre-:*S1pr3^-/-^*;n=13). Each symbol represents one mouse. Scatter dot plots show mean±SEM. Statistical analysis was performed using unpaired t-test and two-way ANOVA followed by Sidak’s, Tukey’s or Bonferroni’s multiple comparison.

Consistent with receptor redundancy in renal vascular resistance, mice with isolated deficiency of S1PR2 in VSMC and cardiomyocytes (*S1pr2*^f/f^ Sm22Cre; Supplemental Fig 4A) or of S1PR3 in all cells (Supplemental Fig 4B) had normal BP. Intriguingly, HRs were increased in S1PR3 deficient mice (Supplemental Fig 4B), suggesting possible compensation for reduced vascular resistance. S1PR2/3 double knockout (dKO) mice die in late gestation with high penetrance (65), and most rare survivors die shortly after weaning with seizures (66). Although we were able to study renal vascular resistance in a few young survivors, high embryonic and postnatal lethality precluded us from measuring BP and vascular resistance in adult dKO mice. Reasoning that genetic compensation was less likely to mask an adult phenotype at the level of ligand than receptor and with postnatal gene deletion, we instead crossed global and conditional *Sphk* knockout alleles into *S1pr2* or *S1pr3* knockout backgrounds and induced *Sphk1* deletion postnatally. Consistent with the dominant role for S1PR3 in S1P-induced renal vascular resistance *ex vivo*, plasma S1P deficiency induced hypotension in S1PR2-deficient males and females (Fig 4F; Supplemental Fig 4C) but had no significant effect on BP in S1PR3-deficient mice of either sex (Fig 4G; Supplemental Fig 4D). Consistent with a tendency in littermate controls (Supplemental Fig 4B), BPs were lower and heart rates higher in S1P proficient controls in the S1PR3 vs. the S1PR2 deficient cohorts (Fig 4F, Supplemental Fig 4,D), arguing that isolated S1PR3 deficiency alone may reduce vascular resistance. Collectively, this argues redundancy between S1PR2&3 with a dominant role for S1PR3 in S1P-mediated regulation of vascular resistance and BP in naïve mice.

### Plasma S1P does not modulate BP responses to dietary salt or Angiotensin II challenge

Our observation that S1P promotes vascular resistance and the spontaneous raise in BP with age may suggest causality in the correlation between circulating S1P and experimental and human hypertension (16, 17, 67). Although our results point to vascular actions of plasma S1P, S1PRs are also expressed in the collective duct and have been implicated in sodium retention (68). To investigate the role of circulating S1P in electrolyte balance and renal and adrenal function, we challenged plasma S1Pless mice and littermate controls with low, normal and high sodium diet (Fig 5A)(64, 69). Consumption of low or high sodium diet for 3 weeks did not impact SBP of littermate controls nor of plasma S1Pless mice, which maintained lower SBP than littermate controls independent of sodium intake (Fig 5B). Consistently, blood sodium and calcium were unchanged, while potassium was lower in plasma S1Pless mice than controls only during low sodium diet (Fig 5C), possibly due to a compensatory decrease in responsiveness to AngII. Physiological parameters including partial (p) O2 or CO2 pressure, HCO3, total (t) CO2, and base excess extracellular fluid (BEecf) were within the normal range and not different between the genotypes (Fig 5D, E), suggesting a normal acid-base status. Urine aldosterone levels increased with low sodium diet and decreased with high sodium diet independent of genotype (Fig 5F). Thus, consistent with a role primarily in vascular tone regulation, plasma S1P is not implicated in renal salt handling nor in adrenal responses to dietary salt.

**Figure 5.**
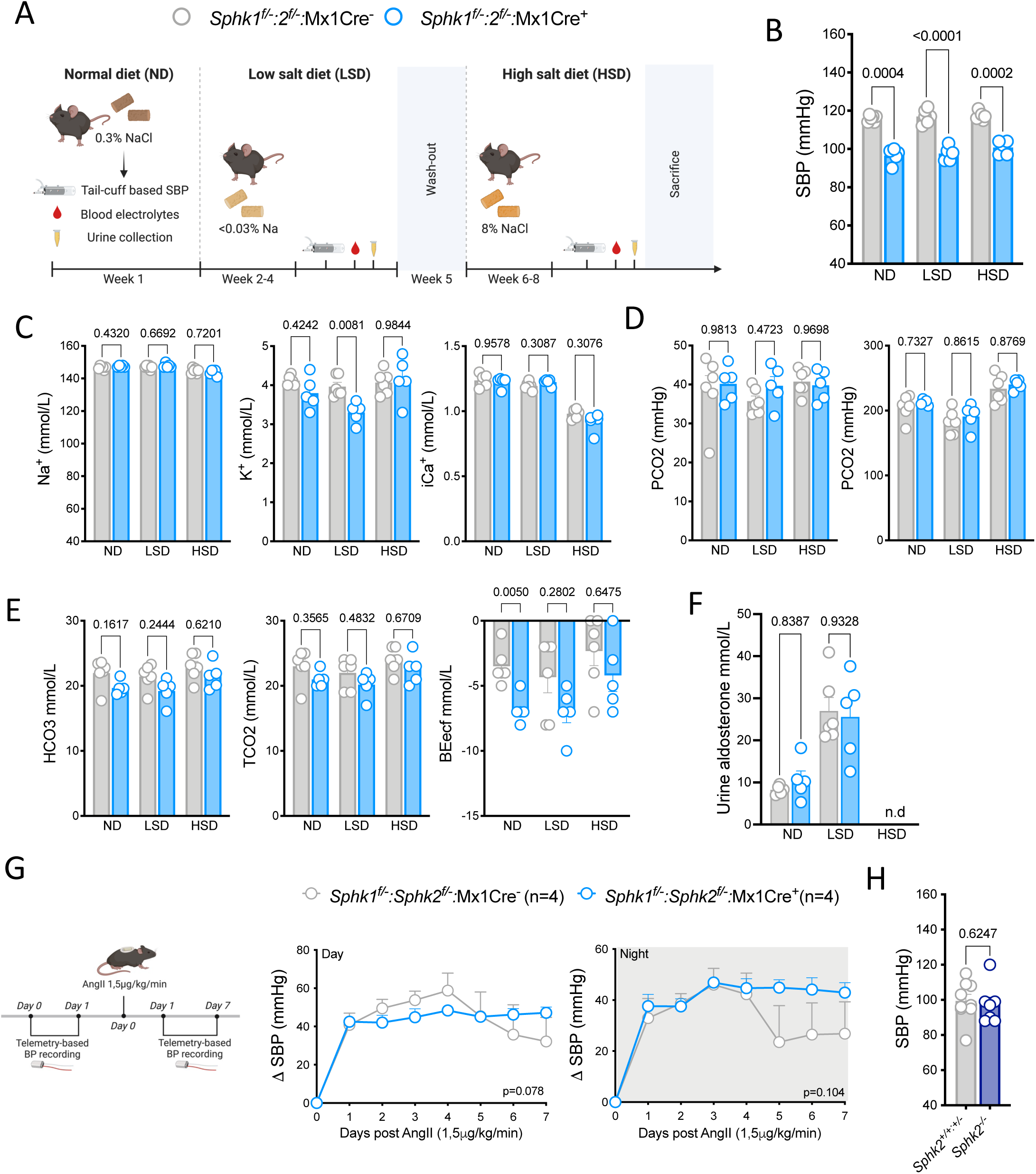
Circulating S1P is dispensable for salt handling and AngII-induced hypertension. **(A)** Experimental design and timeline of dietary salt challenge. (**B**) SBP of plasma S1Pless (*Sphk1*^f/-^:*2*^f/-^: Mx1Cre^-^; n=6) and littermate control (Sphk1^f/-^:2^f/-^: Mx1Cre^+^ ; n=5-6) males measured by tail cuff plethysmography during normal diet (ND), low salt diet (LSD) and high salt diet (HSD). (**C-F**) i-STAT® based blood analysis of ions (Na^+^,K^+^ and iCa^+^) (C), partial pressure of O2 (pO2) and CO2 (pCO2) (D), acid-base balance (bicarbonate (HCO3), total CO2 (TCO2), and base excess extracellular fluid (BEecf) (E) and LC-MS/MS-based urine analysis of aldosterone levels in the same experimental cohort. (**G**) Telemetry based day and night delta SBP of plasma S1Pless (*Sphk1*^f/f^:*2*^-/-^:Mx1Cre^+^; n=4) and littermate control mice (*Sphk1*^f/+^:*2*^-/-^:Mx1Cre^+^; n=4) before and during 7 days of AngII (1.5ug/kg/min) infusion via osmotic minipumps. Left panel shows experimental design and timeline of AngII-induced hypertension. (**H**) Carotid artery SBP measurement of anesthetized Sphk2 deficient (*Sphk2*^-/-^; n=7) and littermate control (*Sphk2*^+/-;+/+^;n=9) males. Each symbol represents one mouse. Scatter dot plots show mean±SEM. Statistical analysis was performed using Mann-Whitney and one-way or two-way ANOVA followed by Bonferroni’s or Sidak’s multiple comparison.

AngII is a key driver of hypertension via actions on multiple organs, including resistance arteries. Sphk1 was identified as an important modulator of vascular function in response to AngII (25) and mice with global Sphk1 or Sphk2 deficiency are both protected from AngII-induced hypertension by a mechanism proposed to involve immune modulation (24). To address if circulating S1P is implicated in AngII-induced hypertension, we implanted telemetry probes into plasma S1Pless mice and littermate controls for measurements of basal pressure and subsequent responses to AngII infusion (500 ng/kg/min) via osmotic minipumps for a period of 4 weeks, during which BP was recorded for 24 hours on day 7, 14, 21 and 28 after infusion (Supplemental Fig 5). The relative increase in SBP with AngII did not differ between genotypes. A similar relative increase in BP was also observed in an acute, high dose AngII model (1500 ng/kg/min) where plasma S1P deficiency was induced in an Sphk2-/-background (*Sphk^f/f^:2^-/-^:*Mx1Cre^+^) (Fig 5G). This argues against a critical role for circulating S1P sources or S1P-dependent lymphocyte egress in AngII-induced hypertension, but does not exclude a role for S1P produced by Mx1Cre-resistent cell types. To address the causal link between plasma S1P levels and pathological increases in BP, we measured BP in anesthetized Sphk2 deficient mice, in which plasma S1P concentrations are twice normal levels due to defective clearance (70). There was no difference in SBP between Sphk2 deficient mice and littermate controls (Fig 5H). Thus, we did not observe a role for plasma S1P in renal salt handling, adrenal responses to changes in dietary salt, nor in AngII-driven hypertension. Moreover, supraphysiological plasma S1P is not sufficient to drive hypertension, and neither Sphk2 nor S1P-mediated lymphocyte trafficking is essential for AngII-induced hypertension.

### Plasma S1P supports cardiac function and reserve

Reduced plasma ApoM and S1P levels and an inverse correlation between left ventricular ejection fraction (LVEF) and S1P levels in HF patients may suggest that plasma S1P also supports cardiac function (14, 15). Despite normal CO (Fig 3B), baseline M-mode echocardiography indeed revealed that LV internal diameters (LVID) were larger in both systole and diastole, and LVEFs reduced in plasma S1Pless males (Fig 6A). As observed with BP probes, HRs were normal (Fig 6A). Septal and posterior wall thicknesses were unchanged (Supplemental Fig 6A), but LV mass slightly increased (Fig 6B), reflecting LV dilation. By contrast, cardiac function and mass in female mice were indistinguishable from littermate controls (Supplemental Fig 6B). Reduced contractile function in male mice was paralleled by modest fibrosis (Supplemental Fig 6C), a slight increase in heart weight to tibia length ratios and mild cardiomyocyte hypertrophy (Supplemental Fig 6D). Thus, basal CO in plasma S1Pless males (Fig 3B) is sustained by compensatory left ventricular (LV) dilation, without major structural alterations to the myocardium. However, a dobutamine stress test revealed reduced cardiac reserve in plasma S1Pless males, characterized by blunted capacity to increase LVEF and CO despite HR elevation similar to controls (Fig 6C). Analysis of Mx1Cre-mediated induction of an mTmG reporter showed little or no activity in VSMC or cardiomyocytes (Supplemental Fig 6E). Accordingly, a similar phenotype of reduced FS, LVEF and increased LVIDs was observed in wild-type recipients of Sphk1&2 deficient BMC (Fig 6D), pointing to a role for S1P derived from hematopoietic cells.

**Figure 6.**
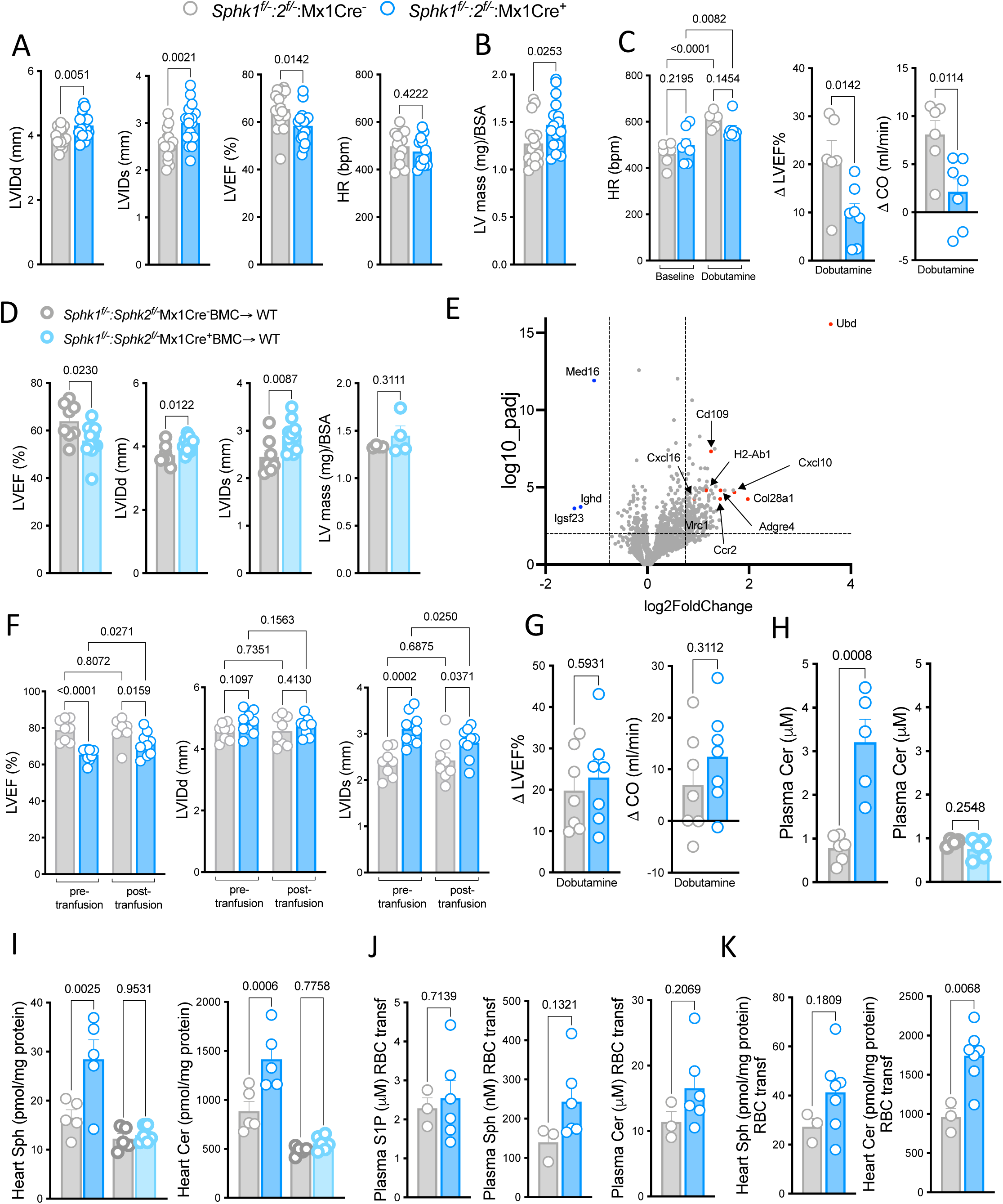
Reduced cardiac function in plasma S1Pless mice. **(A)** Echocardiographic assessment of left ventricular internal diameter in end diastole (LVIDd) and end systole (LVIDs), left ventricle ejection fraction LVEF, heart rate (HR), (**B**) left ventricle mass normalized to body surface area (LV mass/BSA) in plasma S1Pless (*Sphk1*^f/-^:*2*^f/-^:Mx1Cre^+^; n=12-16) and littermate control (*Sphk1*^f/-^:*2*^f/-^:Mx1Cre^_^; n=12-16) males. (**C**) Echocardiographic and Doppler assessment of HR, delta EF and delta CO before and 10 minutes after dobutamine (1,5ug/g, i.p.) administration to *Sphk1^f/-^:Sphk2*^f/-^:Mx1Cre+ (n=7) *Sphk1*^f/-^:*Sphk2*^f/-^:Mx1Cre- (n=6) mice. (**D**) Echocardiographic assessment of LVEF and LV mass/BSA in irradiated wild-type male recipients of *Sphk1*^f/-^:*2*^f/-^: Mx1Cre^+^ (n=12) or *Sphk1*^f/-^:*2*^f/-^:Mx1Cre^-^ (n=8) BMC. (**E**) Volcano plot showing log2 (fold change, FC) against −log10 (p-value) of transcripts identified by RNASeq analysis. Blue dots indicate significant transcripts with a FC ≤ 1.5; red dots indicate transcripts with a FC ≥± 1.5 (**F)** Echocardiographic assessment of LVEF, LVIDd and LVIDs in plasma S1Pless (*Sphk1*^f/-^:*2*^f/-^:Mx1Cre^+^; n=9) and control (*Sphk1*^f/-^:*Sphk2*^f/-^:Mx1Cre^-^; n=8) males pre and 48 hours post erythrocyte transfusion. (**G**) Echocardiographic and Doppler assessment of HR, delta EF and delta CO before and 10 minutes after dobutamine (1,5ug/g, i.p.) administration plasma S1Pless (*Sphk1*^f/-^:*2*^f/-^:Mx1Cre^+^; n=7) and control (*Sphk1*^f/-^:*Sphk2*^f/-^:Mx1Cre^-^; n=7) males 48 hours post erythrocyte transfusion. (**H-J**) Total plasma (H) and heart (I) ceramide levels of plasma S1Pless mice (*Sphk1*^f/-^:*2*^f/-^:Mx1Cre^+^ ; n=5) and littermate controls (*Sphk1*^f/-^:*2*^f/-^:Mx1Cre^+^ ; n=6) with and without erythrocyte transfusion (harvested at 48 hours) and of wild type mice transplanted with BMC from *Sphk1*^f/-^:*2*^f/-^: Mx1Cre+ (n=5-6) or *Sphk1*^f/-^:*2*^f/-^:Mx1Cre^-^ (n=4-5) donors (J). Each symbol represents one mouse. Scatter dot plots show mean±SEM. Statistical analysis was performed using unpaired t-test one and two-way ANOVA followed by Sidak’s, Tukey’s or Bonferroni’s multiple comparison.

To address if the heart undergoes transcriptional reprogramming in the absence of circulating S1P, we performed bulk RNAseq and unbiased gene enrichment analysis on hearts from plasma S1Pless and littermate control males. Of 15341 genes with detectable expression, 1040 were differentially expressed (DEGs), 793 of which were upregulated and 247 downregulated. Of 316 genes with a log2 (fold change (FC))>1,5 nearly all were upregulated (Fig 7E). These were enriched in gene ontology terms for inflammation and macrophages (Supplemental Fig 6F) and dominated by hematopoietic cell transcripts (e.g. cxcl9, cxcl10, cxcl16, ccr2, mpeg1, gpr132, ly86, cd74, itgax, itgal, itga4, cd52, csf2rb, pou2f2, lst1) (Fig 6E). With the exception of 4 genes, the increase in expression was modest (<3 fold). Hallmark genes for fibroblasts/fibrosis and heart failure (e.g. pdgfra, acta2, col22A1, postn, nppb, tnni1, prelp P4ha1, Dcn)(71) were in most cases unchanged, with the exception of the modest increase of some collagens (Col1a1, 1.3 FC, Padj 0.023; col3a1, 1.2 FC, Padj 0.091; col28a1, 3.9 FC, Padj <0.0001), MMPs (mmp12, 2.3 FC, Padj 0.0008; mmp13, 1.7 FC, Padj 0.011; mmp14 1.2 FC, Padj 0.044) and markers of cardiac remodeling (lgas3 2.0 FC, Padj 0.0001). Cross-referencing DEGs to scRNASeq databases and cell marker gene-sets indicated that much of the response could be accounted for by an increase in cardiac macrophages (Supplemental Fig 6F). Increased expression of macrophage markers was confirmed by qPCR and observed also in wild-type recipients of *Sphk1&2* deficient BMC (Supplemental Fig 7A,B). Immunostaining of heart sections from the latter model showed an approximate doubling of mrc1/CD206+ tissue macrophages (Supplemental Fig 7B). Of 15000 genes detected, only ten were expressed at levels <70% of controls, and these did not suggest a specific response. Thus, transcriptional analysis revealed an increase in cardiac macrophages, but did not point to macrophage-induced inflammation nor to transcriptional rewiring of the myocardium.

Basal LV function in 6-8 month old males was significantly improved and cardiac reserve indistinguishable from controls 48 hours after transfusion of wild-type erythrocytes to plasma S1Pless mice (Fig 6F,G), indicating that circulating S1P plays a tonic role in support of cardiac function. Despite this parallel to the rescue of the BP phenotype (Fig 2I), normal basal CO argues that hypotension is not a manifestation of LV dysfunction. Moreover, and not surprisingly given the observed structural changes, LV function was not fully normalized with erythrocyte transfusion (Fig 6F). As might also be expected, macrophage markers remained elevated (Supplemental Fig 7C). Contrasting BP (Fig 4G; Supplemental Fig 4D), the cardiac phenotype persisted in an S1PR3 knockout background (Supplemental Fig 7D). Mice with global S1PR3 deficiency displayed grossly normal cardiac function (Supplemental Fig 7E), as did mice S1PR2 deficiency in VSMC and cardiomyocytes (*S1pr2*^f/f^ Sm22Cre^+^; Supplemental Fig 7F). Mice with neonatally induced EC S1PR1 deficiency, which display a similar vascular leak phenotype to plasma S1Pless mice (19, 28), also had normal LV function and responses to dobutamine (Supplemental Fig 7G). S1PR1 signaling in cardiomyocytes is critical during cardiac development (72), and mice with constitutive deletion of *S1pr1* in cardiomyocytes that survive development also present with progressive cardiomyopathy and compromised responses to dobutamine (73). However, the basal cardiac phenotype of these mice was attributed to embryonic gene deletion (74), and analysis of S1PR1 signaling reporter mice did not show S1PR1 activity in ventricular cardiomyocytes in the mature naïve heart (28, 75). Accordingly, postnatal deletion of S1PR1 in cardiomyocytes did not impact LV systolic function in naïve mice (Supplemental Fig 7H).

Disturbances in the balance of circulating and cardiac sphingolipids may impact cardiac function (27, 76, 77). In accordance with low level Mx1Cre activity in the heart, RNASeq analysis did not reveal changes in the expression of *Sphk1&2* or other enzymes of the sphingolipid pathway to suggest compensatory metabolic rewiring. We have reported that plasma sphingosine is increased in plasma S1Pless mice generated by *Sphk1&2* deletion with Mx1Cre, and that hematopoietic *Sphk1&2* deletion induced with Vav1Cre also results in a slight increase in cardiac sphingosine and ceramide, with no change in cardiac S1P (28). Plasma ceramide was increased in mice with Mx1Cre-mediated *Sphk1&2* deletion, but no significant change was observed wild-type recipients of *Sphk1&2* deficient BMC (Fig 6H), which present a similar cardiac phenotype (Fig 6D). A similar pattern was observed for cardiac sphingosine and ceramide (Fig 6I). Erythrocyte transfusion fully restored plasma S1P levels at 48 hours (suggesting physiological plasma S1P levels at 48h also in the BP rescue experiment, Fig 2I), and partially normalized plasma sphingosine and ceramide levels (Fig 6J). However, cardiac levels of sphingosine and ceramide remained high in these mice (Fig 6K). Thus, although Mx1Cre mediated *Sphk1&2* increases the levels of sphingosine and ceramide in both plasma and cardiac tissue, this increase is not attributable to the hematopoietic compartment and does not track with the cardiac phenotype.

Collectively, our observations reveal a tonic role for circulating S1P in maintaining LV contractile function in mice, but do not identify a non-redundant role for any one S1PR to account for its actions.

## Discussion

We report that circulating S1P plays an essential role in maintaining vascular tone, BP and cardiac function independent of the RAAS. This highlights the relevance of plasma S1P as a biomarker for hypertension (16, 17) and heart failure (14, 15) and points to potential benefit for targeting VSMC S1P receptors for the treatment of microvascular dysfunction and BP normalization. Our observations also suggest that the two main circulating pools of S1P play distinct roles in the optimization of vascular tone and BP. This may be of potential physiological and pathological relevance when there is a local or systemic shift in the balance between HDL-S1P and albumin-S1P, such as during platelet activation and in systemic inflammatory response syndrome (18, 78).

Hypertension is an important contributor to global disease burden (2). Despite an assortment of BP lowering drugs in clinical use, many patients remain hypertensive even when adhering to combination therapy (2, 79, 80). First in line BP lowering drugs include RAAS and calcium channels blockers and thiazide-type diuretics (2). Effective treatment of resistant hypertension is likely to require adjunct targeting of distinct pathways of BP regulation. Often considered a disease of the kidney, there is an increasing appreciation for the role of vascular resistance and GPCR signaling in the development of hypertension (4, 81). Our findings that S1P signaling acts in parallel to the RAAS to regulate BP suggest possible therapeutic potential in targeting this pathway. While S1PR3 appears to play a dominant role in S1P-mediated vascular tone regulation in naïve mice (this study), the contribution of S1PR2 may be greater in individuals with metabolic and cardiovascular disease (21–23, 82), suggesting the need to target both receptors. S1P signaling could potentially also be targeted though ligand supply. While our observations in Sphk2 deficient mice argue that increased plasma S1P may not be sufficient to drive hypertension, higher S1P levels in hypertensive individuals (16, 17) may reflect increased production from a distinct source that could be implicated in BP elevation. Increased Sphk1 expression in the arterial wall observed in experimental models of hypertension would be predicted to trigger unwarranted VSMC S1PR activation (17, 57). Sphk1, S1PR2 and S1PR3 could provide potential targets for BP optimization.

Correction of chronic and acute hypotension also represents an important clinical challenge, for which treatment options are limited. Individuals with low BP are more susceptible to the effects of orthostatic hypotension, where inadequate or delayed constriction of peripheral arteries upon standing results in a temporary reduction of CO and blood flow to the brain. When BP is below normal, cerebral and retinal perfusion can drop so low as to trigger loss of consciousness, blurred vision, confusion and ataxia, resulting in accidental falls. Chronic reductions in cerebral perfusion in orthostatic hypertension may also contribute to ischemic stroke, dementia and cognitive decline (83). When acute and profound, such as after blood loss from traumatic injury, in systemic inflammatory response syndrome and in anaphylactic shock, hypotension can be life endangering. It is notable that levels of albumin-S1P drop sharply in patients with trauma and severe sepsis (18). Inhibitors of S1P lyase, which increase tissue S1P levels, and agonism of S1PR3 were both protective in a murine models of severe sepsis (84). The benefit of albumin infusion for outcome of severe sepsis has been addressed in a number of clinical studies but remains controversial (85). In light of our findings, it may be important to consider how much S1P is bound to the albumin that is infused, and to address the potential benefit of S1P-loaded albumin. In mice, a profound reduction plasma S1P is also observed in systemic anaphylaxis, and infusion of albumin-S1P is protective (19, 34, 37). As a trophic factor for homeostatic maintenance of arterial tone, albumin-S1P deserves further attention as a biomarker and treatment option for hypotension together with agonists for VSMC S1PR3.

S1P circulates in plasma bound to HDL/ApoM and albumin (27). ApoM stabilizes S1P and supports endothelial-dependent vasodilation and vascular integrity through S1PR1 (9, 86, 87). Albumin-S1P on the other hand is turned over rapidly, suggesting that albumin may help shuttle S1P for systemic sphingolipid redistribution and clearance (27, 70). Our observations suggest that albumin-S1P may also play a distinct functional role from HDL-S1P. The distribution of S1P between these two chaperones could also be important for vascular function. When albumin-S1P drops rapidly in trauma and sepsis (18), the relative resistance of HDL-S1P could potentially shift the balance of S1P signaling from S1PR2/3-mediated vasoconstriction to S1PR1-mediated vasodilation, which would be detrimental in this context. It is also notable that the S1P that is released by platelets upon activation associates mainly with albumin (30). If albumin-S1P reaches VSMC S1PRs as indicated in this study, the local increase in S1P concentrations generated upon platelet activation (78) may contribute to vasoconstriction in primary hemostasis (88).

S1P is important for the maintenance of both arterial function and vascular integrity. Suggesting a potential mechanistic link between these functions, we recently reported that S1PR1 activity in the blood vasculature outside of the lung is restricted to resistance-size arteries, and that albumin extravasation is increased in cerebral arterioles from both S1PR1 ECKO mice and plasma S1Pless mice (28) (28). This may suggest that S1P optimizes its own trans-endothelial transport, and presents the possibility that increased albumin extravasation and VSMC S1PR activation may contribute to hypertension observed in S1PR1 ECKO as well as ApoM deficient mice, with also display increased arterial albumin extravasation due to deficient EC S1PR1 activation (10, 89). Such a mechanism could also help explain hypertension reported in patients on fingolimod and other functional S1PR1 antagonists, which has been attributed both to EC S1PR1 desensitization and to S1PR3 activation. Increased transendothelial transport could also be envisioned to indirectly regulate VSMC function by increasing the access of other vasoactive mediators than S1P.

Altered endothelial barrier function does not explain hypotension in plasma S1Pless mice, as the barrier defect but not the BP phenotype is mirrored by EC S1PR1 deficiency (28). Yet, as the endothelium remains a partial barrier to S1P access to VSMC receptors (32), it is conceivable that the increased S1P dependence on BP with age observed in this study relates to age-dependent alterations in barrier function that increase in the exposure of VSMC receptors to albumin-S1P. Indeed, transcellular transport mechanisms that carry albumin across the arterial endothelium increase both in experimental hypertension (90) and aging (91). Moreover, although BP is not increased in naïve Sphk2 deficient mice despite substantially elevated plasma S1P levels, Sphk2 deficiency does not in our hands impair endothelial barrier function. It therefore remains possible that an increase in circulating S1P contributes to increased vascular resistance and blood pressure when endothelial transcytosis is also increased, such as in hypertension (90).

Plasma ApoM and S1P levels correlate inversely with LV function and outcome in heart failure patients, potentially implicating plasma S1P in cardiac function (14, 15). Although not sufficient to impact CO in naïve mice, we observe that disruption of Sphk activity in hematopoietic cells impact both LV function and cardiac reserve, supporting potential causality. Unlike for hypotension, we were not able to attribute this phenotype to a specific deficiency in S1PR signaling. S1PR1 ECKO mice had normal cardiac function even if they display impairment of vascular integrity and hypercapnia-induced hyperemia similar to plasma S1Pless mice (19, 28). This argues against a local or distal effect of endothelial dysfunction. While cardiac function was normal also in mice with deficiency of S1PR1, S1PR2 or S1PR3 in cardiomyocytes alone or together with other cell types, it is notable that roles for cardiomyocyte S1PR2&3 in ischemia/reperfusion injury were only revealed in compound knockouts (92). Receptor redundancy or genetic compensation in cardiomyocytes may therefore have masked a cardiac phenotype in single knockouts the same way redundancy in VSMC masked the BP phenotype. It is also possible that the phenotype involved S1PRs on other cardiac cells or was receptor independent. The ceramide/S1P balance is critical for cardiac function (76, 77), and cardiomyocyte Sphks contribute to heart regeneration (93). Yet the Mx1Cre used to delete *Sphks* in this study did not target cardiomyocytes and the increase in cardiac ceramides and sphingosine observed with hematopoietic Sphk deficiency was modest. Sphk1 also plays an important role in glucose uptake, glycolytic activity and oxygen delivery from erythrocytes (94–96), and it is conceivable that a consequential impairment of oxygen and glucose delivery contribute to the cardiac phenotype of plasma S1Pless mice. While the mechanisms remain to be defined, our study argues that erythrocyte-derived S1P provides trophic support of cardiac function.

By its signaling through S1PR1 on endothelial cells, plasma S1P is known to modulate vascular tone and thus arterial resistance to support blood flow adaptation to local metabolic needs by engaging eNOS and NO-mediated vasodilation. By engaging arteriolar VSMCs through S1PR2&3 and cardiomyocytes through mechanisms yet to be defined, our study suggests that plasma S1P also optimizes the levels of total arterial resistance and left ventricular systolic function to maintain cardiovascular homeostasis and BP. Different pools of chaperone-associated S1P appear to perform these actions, with HDL-associated S1P mainly vasodilatory and albumin-S1P mainly contractile, suggesting diagnostic benefit of differentiating between these plasma pools and the importance of analyzing plasma rather than serum to avoid contamination from platelet S1P.

## Funding

This work was supported by the French National Research Agency (ANR-19-CE14-0028, ANR-21-CE17-0023, EC,DH,PB); the program “Investissement d’Avenir” launched by the French Government and implemented by ANR (ANR-18-IdEx-0001) as part of its program « Emergence », the French Foundation for Medical Research (DCP20171138945; EC); Singapore Ministry of Health’s National Research Council (NMRC/OFIRG/0066/20, LNN), Singapore Ministry of Education (MOE2018-T2-1-126, T2EP30123-0014 and NUHSRO/2022/067/T1; LNN.); Fondation de France (EC), the French society for arterial hypertension (BM), the Lefoulon-Delalande Foundation (IDG), France Génomique (ANR-10-INBS-09; the Institut Pasteur Biomics Platform) and IBISA (the Institut Pasteur Biomics Platform).

## Acknowledgements

We are very grateful for support from the technical and administrative platforms at the Paris Cardiovascular Research Center. We would like to thank Ivo Cornelissen, Tovo David, Kathy Ruppel, Shaun Coughlin, Stephen Wilson, Daniel Duong, Salome Gazit and Maria Chavez Canales for help with this project and Sonia Bergaya, Juliette Hadchouel, Sheerazed Boulkroun and Neeraj Dhaun for helpful discussions and advice. The graphical abstract was created with BioRender.com. We would also like to thank Elodie Turc for help with RNA QC, library generation, sequencing, QC sequencing and Laure Lemée and Etienne Kornobis for help with data management and analysis of RNASeq results.

## Author contributions

IDG performed immunohistochemistry, microscopy and image analysis and tail cuff-based BP measurements, IDG and LC tissue perfusion, organ harvest, and mouse colony management, PB and AB Echo Doppler measurements and analysis, MG, CP and EJ-V renal perfusion and data analysis, VB and ER invasive BP and HR measurements, NM, NF and AN immunofluorescence staining, SB aldosterone measurements, HTTH lipidomic analysis, JC, PLT, OL provided reagents and scientific input, IDG, EJ-V, PB, MCZ, LNN, DH and EC designed experiments and analyzed data, IDG and EC prepared the figures and wrote the paper.

## Competing interests

None to disclose.

---

**Supplemental Figure 1.**
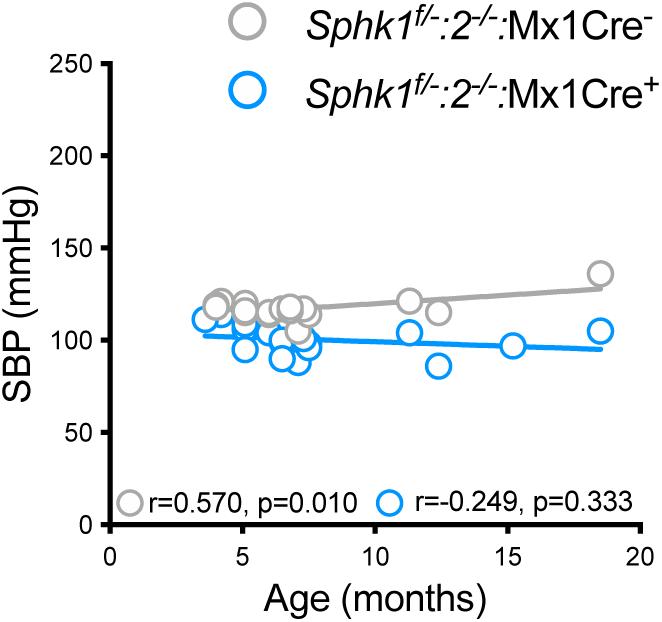
Plasma S1Pless mice are protected from age-induced hypertension. Pearson correlation between age and diurnal SBP measured by non-invasive tail cuff plethysmography for plasma S1Pless (*Sphk1*^f/-^:*Sphk2*^f/-^:Mx1Cre^+^; n=17) and for littermate control (*Sphk1*^f/-^:*Sphk2* ^f/-^:Mx1Cre-(n=19) males. Each symbol represents one mouse. Pearson r and p values displayed.

**Supplemental Figure 2.**
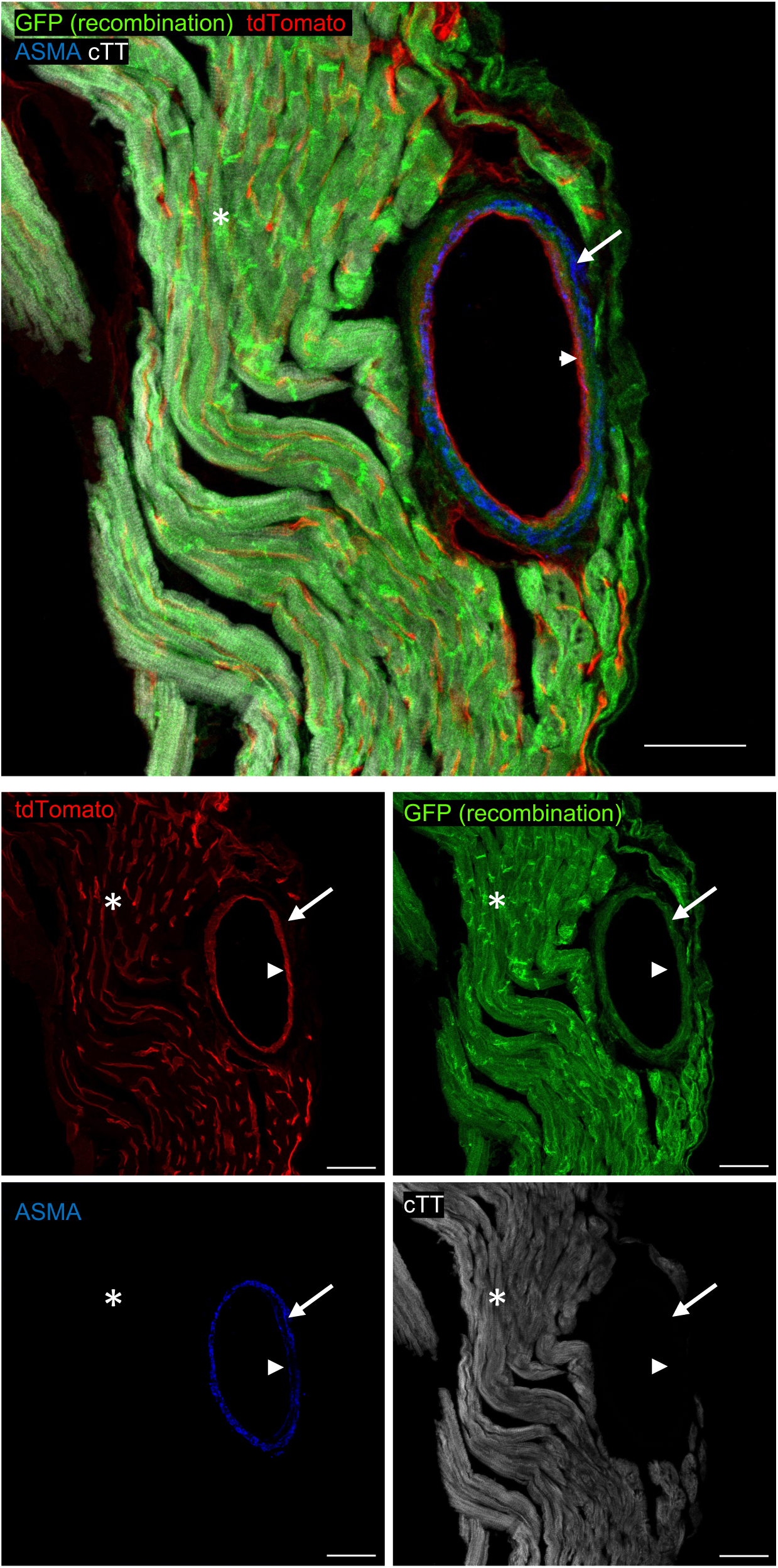
Sm22Cre-induced recombination in the heart. Efficacy and specificity Sm22Cre-induced recombination was assessed with a ROSA^mT/mG^ reporter in which expression of a cell membrane localized red fluorescent protein (tdTomato/mT) is replaced with expression of a cell membrane localized green fluorescent protein (EGFP/mG) upon Cre-mediated recombination of a loxP-flanked stop codon. Heart sections were counter stained with antibodies against ASMA and cardiac troponin T to additionally label VSMCs (blue) and cardiomyocytes (white), respectively. Note homogeneous recombination in both VSMCs (arrow) and cardiomyocytes (asterisk) but not in the endothelium (arrowhead). Images are representative of sections stained from 3 mice. Scale bar: 50 µm.

**Supplemental Figure 3.**
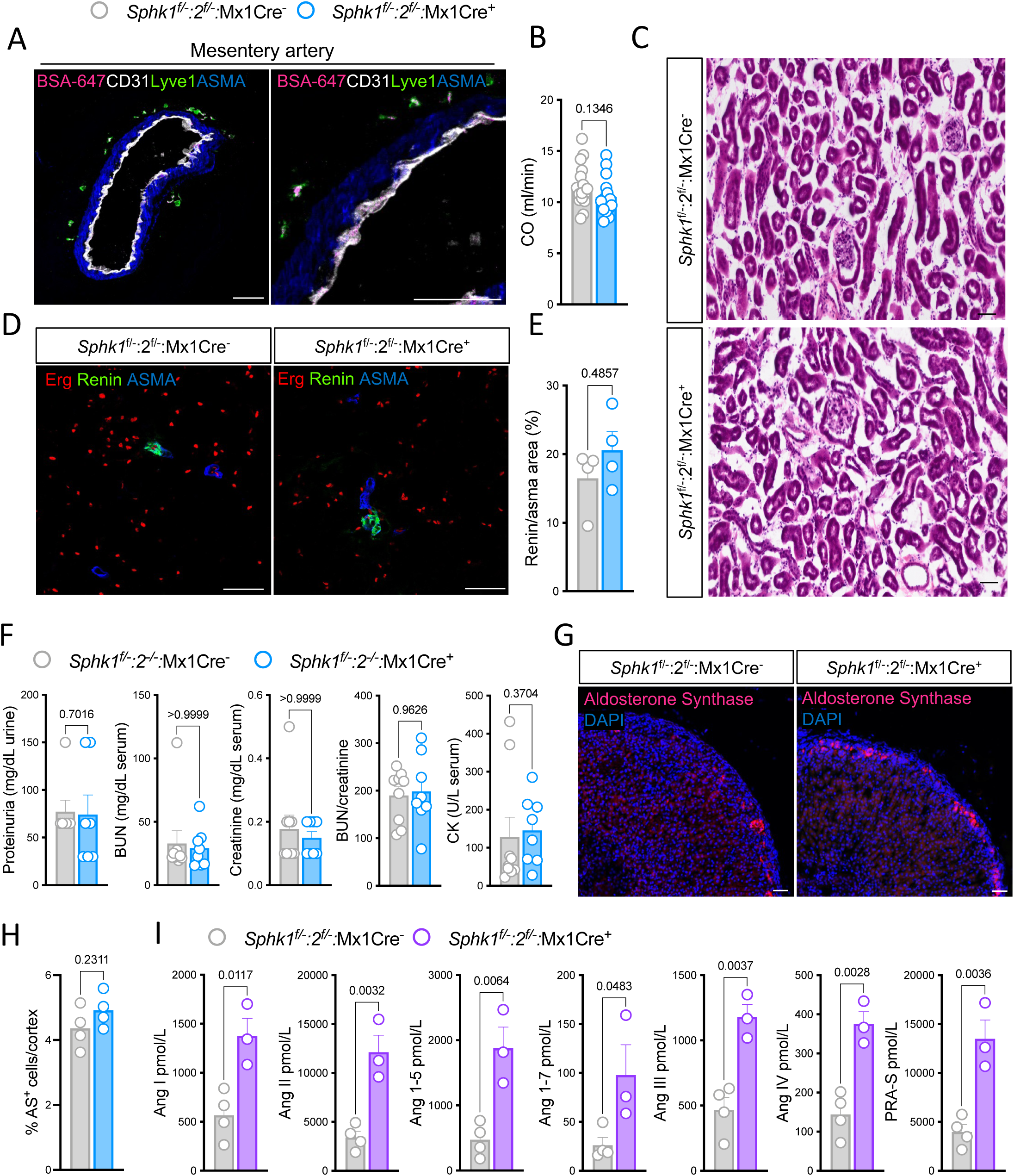
Renal and adrenal function are not compromised by plasma S1P deficiency. (**A**) Accumulation of AlexaFluor647-conjugated BSA (magenta) in mesenteric artery from wild type mice 1 hour after i.v tracer injection counter-stained with antibodies to CD31 to identify EC (white), Lyve1 to identify macrophages (green) and ASMA to identify VSMC (blue). Scale bar 50µm. Right panel shows magnification of the wall of the vessels shown in the left panel. Image is representative of arteries harvested from 3 mice. (**B**) M-mode based echocardiography of cardiac output of plasma S1Pless (*Sphk1*^f/-^:*Sphk2*^f/-^:Mx1Cre^+^;n=16) and littermate control mice (*Sphk1*^f/-^:*Sphk2*^f/-^:Mx1Cre^-^; n=15). (**C)** Hematoxylin and Eosin (H&E) stained kidney sections of plasma S1Pless (*Sphk1*^f/-^:*Sphk2*^f/-^:Mx1Cre^+^) and littermate control (*Sphk1*^f/-^:*Sphk2*^f/-^:Mx1Cre^-^) males. Images are from the renal cortex and representative of n=4 mice for each genotype (**D, E**) Assessment of renin expression in kidney sections of plasma S1Pless (*Sphk1*^f/-^:*Sphk2*^f/-^:Mx1Cre^+^) and littermate control (*Sphk1*^f/-^:*Sphk2*^f/-^:Mx1Cre^-^) males D, representative images of sections stained for Erg+ EC nuclei (red), renin (green) and ASMA (bue). E, quantification of renin on ASMA occupied area (n=4). (**F**) Assessment of kidney function through measurement of proteinuria (n=7), serum blood urea nitrogen (BUN), creatinine, BUN/creatinine ration and creatine kinase (CK) in plasma S1Pless (*Sphk1*^f/f^:*Sphk2*^-/-^:Mx1Cre^+^;n=7) and control (*Sphk1*^f/f^:*Sphk2*^-/-^:Mx1Cre^-^;n=9) mice. (**G, H**) Confocal microscopy images of sections of adrenal cortex from plasma S1Pless (*Sphk1*^f/f^:*Sphk2*^-/-^:Mx1Cre^+^;n=4) and control (*Sphk1*^f/f^:*Sphk2*^-/-^:Mx1Cre^-^;n=4) mice stained for aldosterone synthase (AS; magenta) and cell nuclei (DAPI, blue). F, Representative images. G, quantification. (**I**) Equilibrium angiotensin metabolite profile of plasma S1Pless (*Sphk1*^f/-^:*Sphk2*^f/-^:Mx1Cre^+^; n=3) and littermate control (*Sphk1*^f/-^:*Sphk2*^f/-^:Mx1Cre^-^;n=4) females. Renin activity (PRA-S) is calculated as the sum of AngI+AngII, and ACE activity (ACE-S) as ratio between Ang II and Ang I. Each symbol represents one mouse. Scatter dot plots show mean±SEM. Statistical analysis was performed using an unpaired t-test.

**Supplemental Figure 4.**
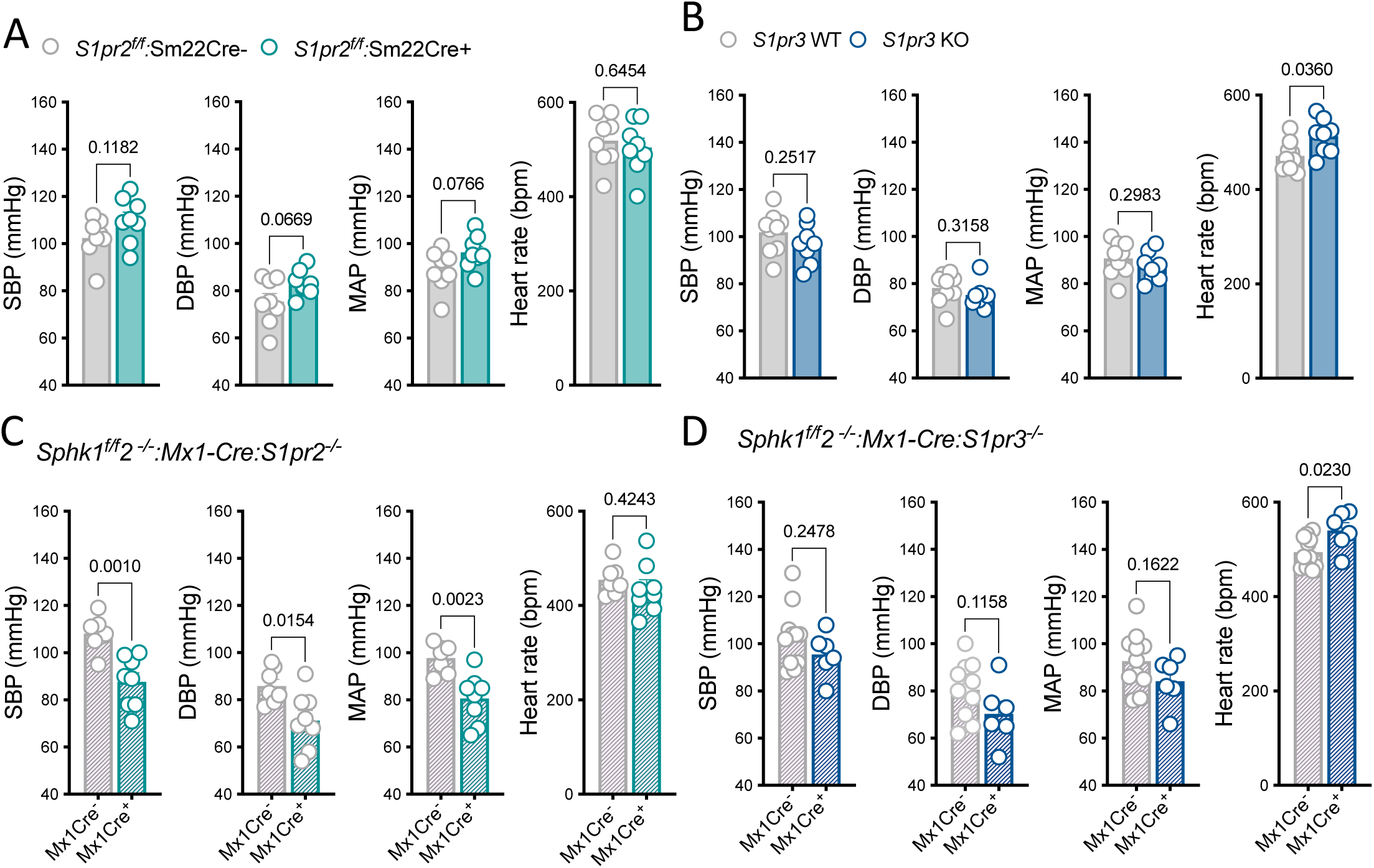
S1PR3 acts downstream of S1P to control blood pressure. (**A, B**) Invasive central blood pressure measurements in anesthetized male mice lacking S1PR2 in cardiomyocytes and VSMC (*S1pr2^f/f^*:Sm22 ; n=8; A) or S1pr3 in all cells (S1pr3^-/-^ n=8; B) and their respective littermate controls (n=8-9). (**C, D**) Invasive central blood pressure measurements in anesthetized female S1P deficient mice and littermate controls in the context of S1PR2 deficiency (*Sphk1*^f/f^:*Sphk2^-/-^:*Mx1Cre^+^:*S1pr2^-/-^* (n=8) vs. *Sphk1*^f/f^:*Sphk2*^-/-^:Mx1Cre^-^:*S1pr2^-/-^* (n=7))(C) or in the context of S1P3 deficiency (*Sphk1*^f/f^:*Sphk2*^-/-^:Mx1Cre^+^:*S1pr3^-/-^* (n=6) vs. *Sphk1*^f/f^:*Sphk2^-/-^*: Mx1Cre^-^:*S1pr3^-/-^* (n=11),)(D). Each symbol represents one mouse. Scatter dot plots show mean±SEM. Statistical analysis was performed using an unpaired t-test.

**Supplemental Figure 5.**
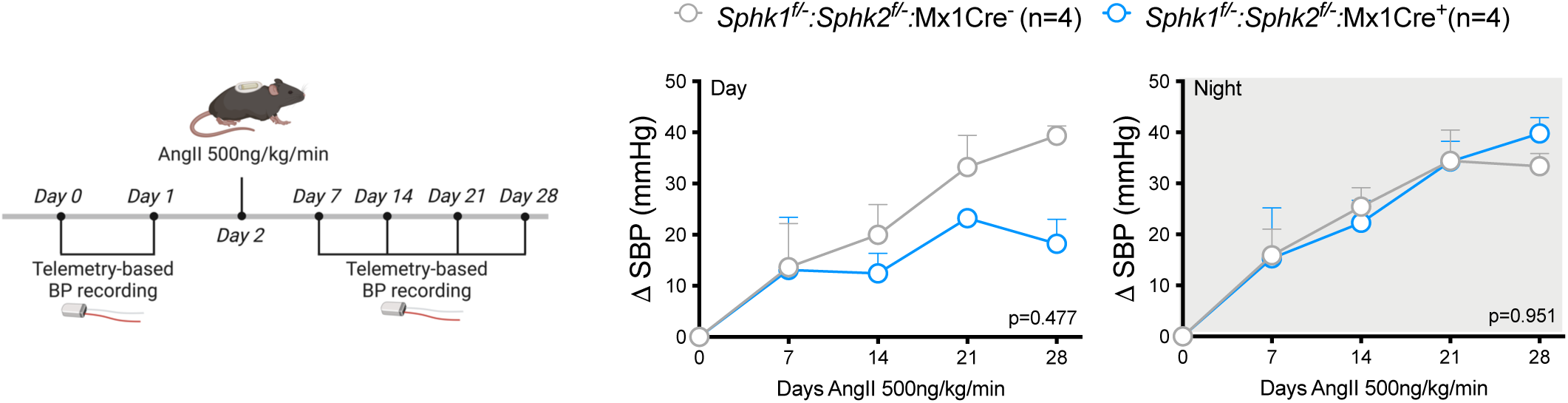
Plasma S1P deficiency does not prevent the development of hypertension during AngII infusion. Telemetry-based night and day SBP measurements of mice (*Sphk1*^f/-^:*Sphk2*^f/-^:Mx1Cre^+^; n=4) and littermate controls (*Sphk1*^f/-^:*Sphk2*^f/-^:Mx1Cre^-^; n=4) recorded weekly during 28 days of AngII (500ng/kg/min) infusion via osmotic minipumps, expressed relative to average pre-perfusion SBPs. Graphs show mean±SEM. Statistical analysis was performed using two-way ANOVA followed by Sidak’s multiple comparison test.

**Supplemental Figure 6.**
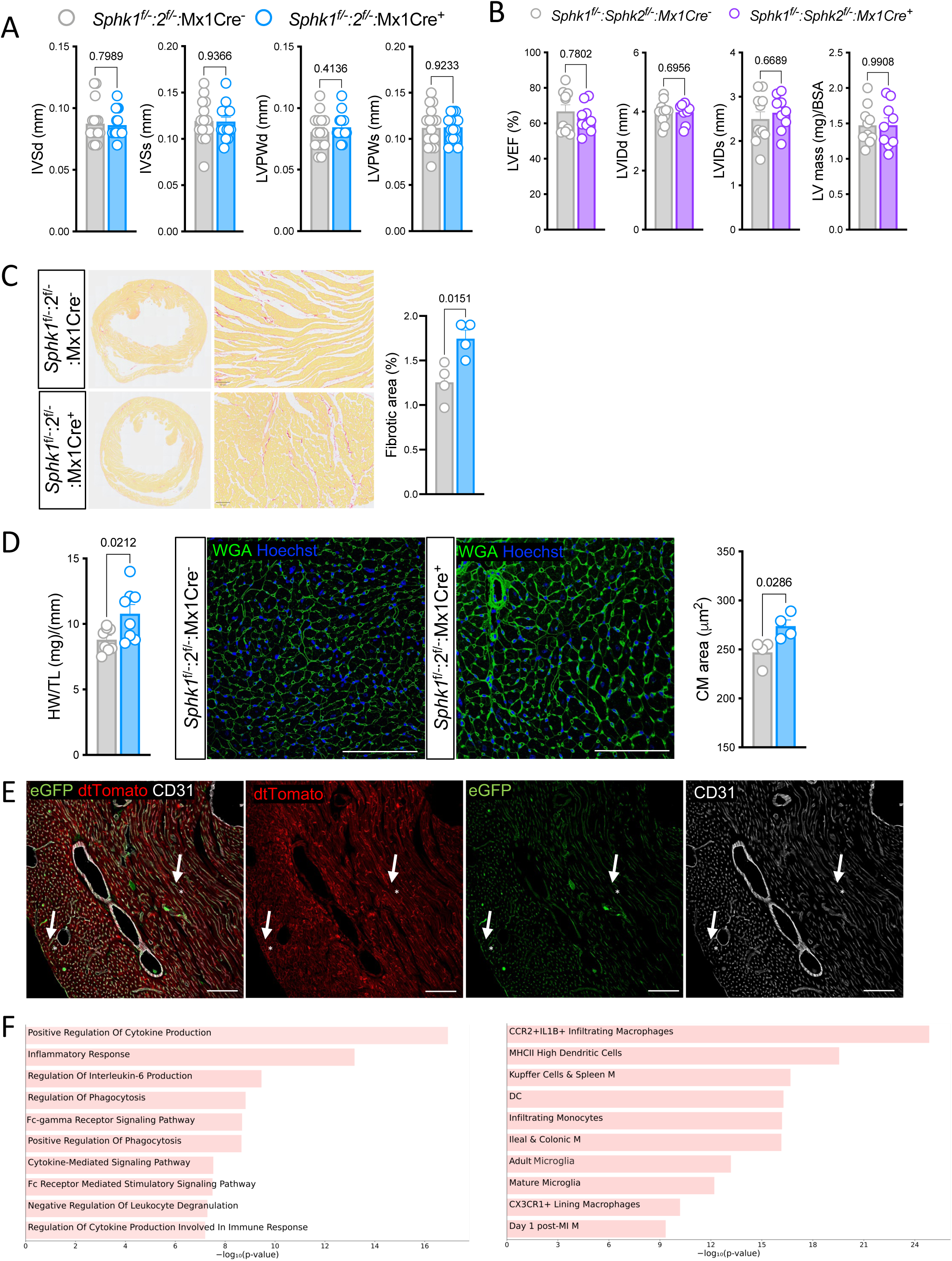
Loss of plasma S1P is associated with mild cardiac remodelling. (**A**) Echocardiographic measurements of interventricular septal end diastolic diameter (IVSd), end systolic diameter (IVSs), left ventricular posterior wall end diastolic diameter (LVPWd) and end systolic diameter (LVPWs) of plasma S1Pless (*Sphk1*^f/-^:*Sphk2*^f/-^:Mx1Cre^+^; n=16) and littermate control (*Sphk1*^f/-^:*Sphk2*^f/-^:Mx1Cre^-^; n=15) males. (**B**) Echocardiographic measurements of left ventricular internal diameter in end diastole (LVIDd) and end systole (LVIDs), left ventricle ejection fraction (LVEF), and left ventricle mass normalized to body surface area (LV mass/BSA) in plasma S1Pless (*Sphk1*^f/-^:*Sphk2*^f/-^:Mx1Cre^+^; n=9) and littermate control (*Sphk1*^f/-^:*Sphk2*^f/-^:Mx1Cre^-^; n=10) females. (**C)** Assessment of cardiac fibrosis in heart sections plasma S1Pless (*Sphk1*^f/-^:*Sphk2*^f/-^:Mx1Cre^+^; n=4) and littermate control (*Sphk1*^f/-^:*Sphk2*^f/-^:Mx1Cre^-^; n=4) males by picosirius red staining. Left panel: representative images hearts; Right panel: quantification of fibrotic area. (**D)** left panel: Quantitative analysis of heart weight/tibia length in plasma S1Pless (*Sphk1*^f/-^:*Sphk2*^f/-^:Mx1Cre^+^; n=8) and littermate control (*Sphk1*^f/-^:*Sphk2*^f/-^:Mx1Cre^-^; n=8) males. Middle panel: Representative immunofluorescent images of heart sections from *Sphk1*^f/-^:*Sphk2*^f/-^:Mx1Cre+ mice and littermate controls (n=4) stained with FITC-labelled wheat germ agglutinin (WGA) and Hoechst; Right panel: quantification of cardiomyocyte (CM) cross-sectional area. (**E)** Excision specificity and efficiency of the Mx1Cre driver was assessed with a ROSA^mTmG^ reporter in adult mice using the same neonatal induction protocol used to generate plasma S1Pless mice. Heart sections were counter-stained with antibodies against CD31 to label EC (white). Note Mx1Cre induced excision (green cells) in EC (arrowhead) but not in vascular smooth muscle cells (arrow) or cardiomyocytes (arrow). Images are representative of sections from 3 mice. Scale bar: 100 µm. **F**, Enrichr based gene set enrichment analysis showing biological processes (left) and cell types (right) enriched in hearts from *Sphk1*^f/-^:*Sphk2*^f/-^:Mx1Cre+ mice compared to littermate controls

**Supplemental Figure 7.**
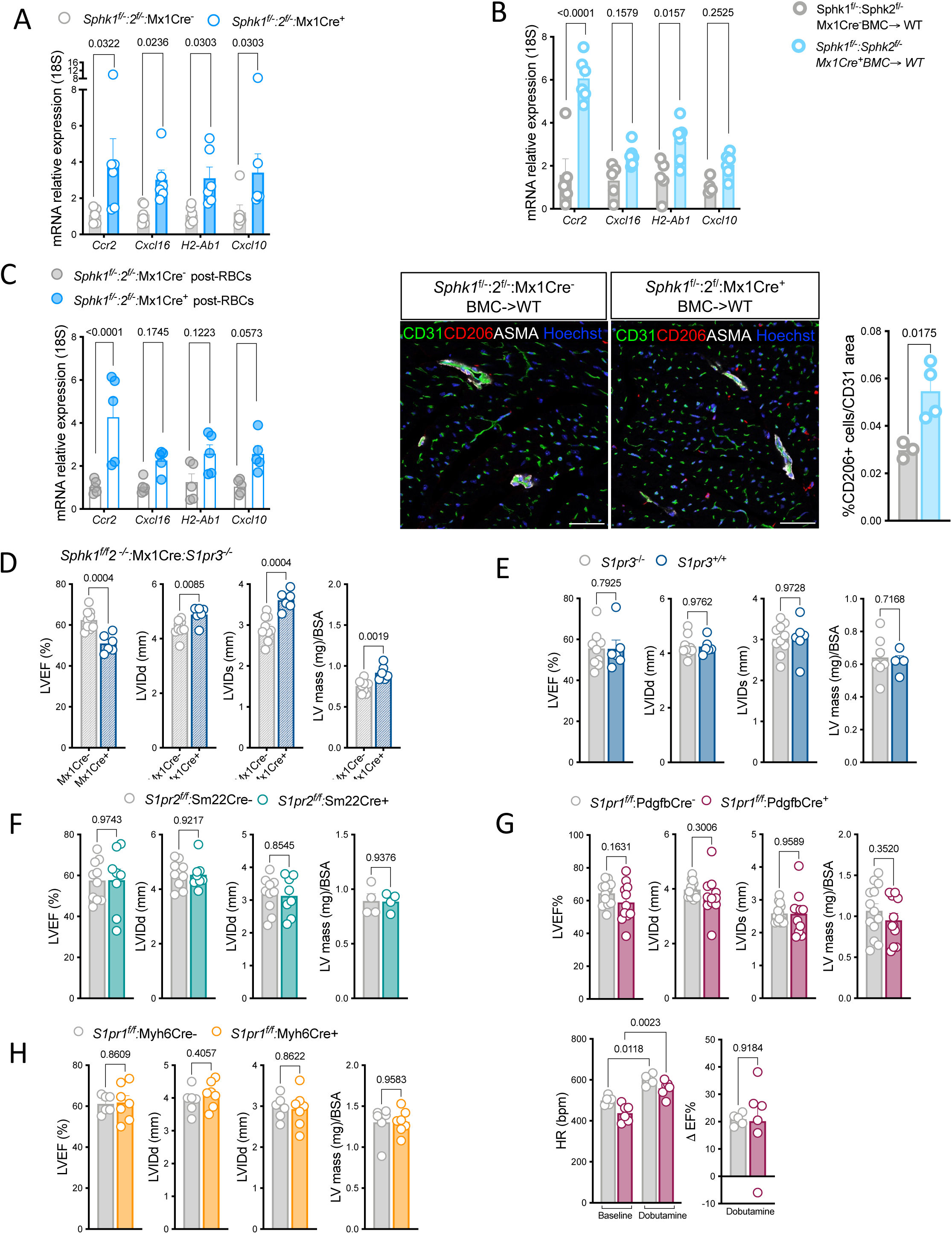
Assessment of mechanisms driving LV dysfunction in plasma S1Pless mice. **(A-C**) Relative mRNA expression of macrophage markers in the hearts of plasma S1Pless (*Sphk1*^f/-^:*Sphk2*^f/-^:Mx1Cre^+^; n=) and control (*Sphk1*^f/-^:*Sphk2*^f/-^:Mx1Cre^-^; n=) males; in lethally irradiated wild-type male recipients of S1Pless (*Sphk1*^f/-^:*Sphk2*^f/-^:Mx1Cre^+^; n=6) and control (*Sphk1*^f/-^:*Sphk2*^f/-^:Mx1Cre^-^; n=5) bone marrow cells (BMC)(B); and in plasma S1Pless (*Sphk1*^f/-^:*Sphk2*^f/-^:Mx1Cre^+^; n=) and control (*Sphk1*^f/-^:*Sphk2*^f/-^:Mx1Cre^-^; n=) males 48 hours after transfusion of erythrocytes from wild-type donors (C). (B), lower panel: Immunostaining for macrophages in heart sections of in lethally irradiated wild-type male recipients of S1Pless (*Sphk1*^f/-^:*Sphk2*^f/-^:Mx1Cre^+^; n=4) and control (*Sphk1*^f/-^:*Sphk2*^f/-^:Mx1Cre^-^; n=4) BMC. Representative images of sections immune-stained with macrophage (CD206), endothelial cell (CD31) and vascular smooth muscle cell (ASMA) markers and counter-stained for cell nuclei (Hoechst) on the left. Scale bar:50 µm. Quantification of CD206+ tissue macrophages (as % of heart tissue) on the right. (**D-H**) Echocardiographic assessment of left ventricular internal diameter in end diastole (LVIDd) and end systole (LVIDs), left ventricle ejection fraction (LVEF) and left ventricle mass normalized to body surface area (LV mass/BSA) of plasma S1Pless (*Sphk1^f/f^:Sphk2^-/-^:*Mx1Cre^+^:*S1pr3^-/-^*; n=6) and littermate control (*Sphk1^f/f^:Sphk2^-/-^*:Mx1Cre^-^:*S1pr3^-/-^*; n=9) males in a background of S1PR3 deficiency (D); globally S1PR3 deficient (*S1pr3^-/-^*; n=6) and littermate control (*S1pr3^+/-;+/+^*; n=9) males; (E), constitutive myocyte S1PR2 deficient (*S1pr2*^f/f^ SM22Cre^+^; n=8) and littermate control (*S1pr2*^f/f^ Sm22Cre^-^; n=10) males (F); neonatally induced endothelial S1PR1 deficient (*S1pr1*^f/f^:PdgfbiCreERT2Cre^+^ ; n=11) and littermate control (*S1pr1*^f/f^:PdgfbiCreERT2Cre^-^ ; n=13) males (G); and neonatally induced cardiomyocyte S1PR1 deficient (*S1pr1*^f/f^:Myh6CreERT2Cre^+^ ; n=7) and littermate control (*S1pr1*^f/f^:Myh6CreERT2Cre^-^ ; n=6) males (H). For endothelial S1PR1 deficient males, HR and delta EF are also shown after stress testing with dobutamine (Cre+ n=6; Cre-n=5) mice (G). Each symbol represents one mouse. Scatter dot plots show mean±SEM. Statistical analysis was performed using unpaired t-test or two-way ANOVA (dobutamine stress test in G).

## Notes

### Competing Interest Statement

The authors have declared no competing interest.

## References

1. Deanfield JE, Halcox JP, and Rabelink TJ. Endothelial function and dysfunction: testing and clinical relevance. Circulation. 2007;115(10):1285–95.

2. Brouwers S, Sudano I, Kokubo Y, and Sulaica EM. Arterial hypertension. Lancet. 2021;398(10296):249–61.

3. Wettschureck N, and Offermanns S. Mammalian G proteins and their cell type specific functions. Physiol Rev. 2005;85(4):1159–204.

4. Althoff TF, and Offermanns S. G-protein-mediated signaling in vascular smooth muscle cells - implications for vascular disease. J Mol Med (Berl). 2015;93(9):973–81.

5. Wirth A, Benyo Z, Lukasova M, Leutgeb B, Wettschureck N, Gorbey S, et al. G12-G13-LARG-mediated signaling in vascular smooth muscle is required for salt-induced hypertension. Nature medicine. 2008;14(1):64–8.

6. Wirth A, Wang S, Takefuji M, Tang C, Althoff TF, Schweda F, et al. Age-dependent blood pressure elevation is due to increased vascular smooth muscle tone mediated by G-protein signalling. Cardiovascular research. 2016;109(1):131–40.

7. Wang S, Iring A, Strilic B, Albarran Juarez J, Kaur H, Troidl K, et al. P2Y(2) and Gq/G(1)(1) control blood pressure by mediating endothelial mechanotransduction. The Journal of clinical investigation. 2015;125(8):3077–86.

8. Igarashi J, and Michel T. Sphingosine-1-phosphate and modulation of vascular tone. Cardiovascular research. 2009;82(2):212–20.

9. Levkau B. HDL-S1P: cardiovascular functions, disease-associated alterations, and therapeutic applications. Front Pharmacol. 2015;6:243.

10. Cantalupo A, Gargiulo A, Dautaj E, Liu C, Zhang Y, Hla T, et al. S1PR1 (Sphingosine-1-Phosphate Receptor 1) Signaling Regulates Blood Flow and Pressure. Hypertension. 2017;70(2):426–34.

11. Nitzsche A, Poittevin M, Benarab A, Bonnin P, Faraco G, Uchida H, et al. Endothelial S1P1 Signaling Counteracts Infarct Expansion in Ischemic Stroke. Circulation research. 2021;128(3):363–82.

12. Christoffersen C, Obinata H, Kumaraswamy SB, Galvani S, Ahnstrom J, Sevvana M, et al. Endothelium-protective sphingosine-1-phosphate provided by HDL-associated apolipoprotein M. Proceedings of the National Academy of Sciences of the United States of America. 2011;108(23):9613–8.

13. Swendeman SL, Xiong Y, Cantalupo A, Yuan H, Burg N, Hisano Y, et al. An engineered S1P chaperone attenuates hypertension and ischemic injury. Science signaling. 2017;10(492).

14. Polzin A, Piayda K, Keul P, Dannenberg L, Mohring A, Graler M, et al. Plasma sphingosine-1-phosphate concentrations are associated with systolic heart failure in patients with ischemic heart disease. J Mol Cell Cardiol. 2017;110:35–7.

15. Chirinos JA, Zhao L, Jia Y, Frej C, Adamo L, Mann D, et al. Reduced Apolipoprotein M and Adverse Outcomes Across the Spectrum of Human Heart Failure. Circulation. 2020;141(18):1463–76.

16. Jujic A, Matthes F, Vanherle L, Petzka H, Orho-Melander M, Nilsson PM, et al. Plasma S1P (Sphingosine-1-Phosphate) Links to Hypertension and Biomarkers of Inflammation and Cardiovascular Disease: Findings From a Translational Investigation. Hypertension. 2021;78(1):195–209.

17. Di Pietro P, Carrizzo A, Sommella E, Oliveti M, Iacoviello L, Di Castelnuovo A, et al. Targeting the ASMase/S1P pathway protects from sortilin-evoked vascular damage in hypertension. The Journal of clinical investigation. 2022;132(3).

18. Winkler MS, Martz KB, Nierhaus A, Daum G, Schwedhelm E, Kluge S, et al. Loss of sphingosine 1-phosphate (S1P) in septic shock is predominantly caused by decreased levels of high-density lipoproteins (HDL). J Intensive Care. 2019;7:23.

19. Gazit SL, Mariko B, Therond P, Decouture B, Xiong Y, Couty L, et al. Platelet and Erythrocyte Sources of S1P Are Redundant for Vascular Development and Homeostasis, but Both Rendered Essential After Plasma S1P Depletion in Anaphylactic Shock. Circulation research. 2016;119(8):e110–26.

20. Lorenz JN, Arend LJ, Robitz R, Paul RJ, and MacLennan AJ. Vascular dysfunction in S1P2 sphingosine 1-phosphate receptor knockout mice. Am J Physiol Regul Integr Comp Physiol. 2007;292(1):R440–6.

21. Sauve M, Hui SK, Dinh DD, Foltz WD, Momen A, Nedospasov SA, et al. Tumor Necrosis Factor/Sphingosine-1-Phosphate Signaling Augments Resistance Artery Myogenic Tone in Diabetes. Diabetes. 2016;65(7):1916–28.

22. Kroetsch JT, and Bolz SS. The TNF-alpha/sphingosine-1-phosphate signaling axis drives myogenic responsiveness in heart failure. J Vasc Res. 2013;50(3):177–85.

23. Hoefer J, Azam MA, Kroetsch JT, Leong-Poi H, Momen MA, Voigtlaender-Bolz J, et al. Sphingosine-1-phosphate-dependent activation of p38 MAPK maintains elevated peripheral resistance in heart failure through increased myogenic vasoconstriction. Circulation research. 2010;107(7):923–33.

24. Meissner A, Miro F, Jimenez-Altayo F, Jurado A, Vila E, and Planas AM. Sphingosine-1-phosphate signalling-a key player in the pathogenesis of Angiotensin II-induced hypertension. Cardiovascular research. 2017;113(2):123–33.

25. Jozefczuk E, Nosalski R, Saju B, Crespo E, Szczepaniak P, Guzik TJ, et al. Cardiovascular Effects of Pharmacological Targeting of Sphingosine Kinase 1. Hypertension. 2020;75(2):383–92.

26. Chen J, Tang H, Sysol JR, Moreno-Vinasco L, Shioura KM, Chen T, et al. The sphingosine kinase 1/sphingosine-1-phosphate pathway in pulmonary arterial hypertension. Am J Respir Crit Care Med. 2014;190(9):1032–43.

27. Kuo A, and Hla T. Regulation of cellular and systemic sphingolipid homeostasis. Nat Rev Mol Cell Biol. 2024.

28. Del Gaudio I, Nitzsche A, Boye K, Bonnin P, Poulet M, Nguyen TQ, et al. Zonation and ligand and dose dependence of sphingosine 1-phosphate receptor-1 signalling in blood and lymphatic vasculature. Cardiovascular research. 2024;120(14):1794–810.

29. Pappu R, Schwab SR, Cornelissen I, Pereira JP, Regard JB, Xu Y, et al. Promotion of lymphocyte egress into blood and lymph by distinct sources of sphingosine-1-phosphate. Science. 2007;316(5822):295–8.

30. Frej C, Andersson A, Larsson B, Guo LJ, Norstrom E, Happonen KE, et al. Quantification of sphingosine 1-phosphate by validated LC-MS/MS method revealing strong correlation with apolipoprotein M in plasma but not in serum due to platelet activation during blood coagulation. Anal Bioanal Chem. 2015;407(28):8533–42.

31. Jang E, Robert J, Rohrer L, von Eckardstein A, and Lee WL. Transendothelial transport of lipoproteins. Atherosclerosis. 2020;315:111–25.

32. Kerage D, Gombos RB, Wang S, Brown M, and Hemmings DG. Sphingosine 1-phosphate-induced nitric oxide production simultaneously controls endothelial barrier function and vascular tone in resistance arteries. Vascular pharmacology. 2021;140:106874.

33. Wafa D, Koch N, Kovacs J, Kerek M, Proia RL, Tigyi GJ, et al. Opposing Roles of S1P(3) Receptors in Myocardial Function. Cells. 2020;9(8).

34. Olivera A, Eisner C, Kitamura Y, Dillahunt S, Allende L, Tuymetova G, et al. Sphingosine kinase 1 and sphingosine-1-phosphate receptor 2 are vital to recovery from anaphylactic shock in mice. The Journal of clinical investigation. 2010;120(5):1429–40.

35. Camerer E, Regard JB, Cornelissen I, Srinivasan Y, Duong DN, Palmer D, et al. Sphingosine-1-phosphate in the plasma compartment regulates basal and inflammation-induced vascular leak in mice. The Journal of clinical investigation. 2009;119(7):1871–9.

36. Vu TM, Ishizu AN, Foo JC, Toh XR, Zhang F, Whee DM, et al. Mfsd2b is essential for the sphingosine-1-phosphate export in erythrocytes and platelets. Nature. 2017;550(7677):524–8.

37. Le TNU, Nguyen TQ, Kalailingam P, Nguyen YTK, Sukumar VK, Tan CKH, et al. Mfsd2b and Spns2 are essential for maintenance of blood vessels during development and in anaphylactic shock. Cell Rep. 2022;40(7):111208.

38. Goppner C, Orozco IJ, Hoegg-Beiler MB, Soria AH, Hubner CA, Fernandes-Rosa FL, et al. Pathogenesis of hypertension in a mouse model for human CLCN2 related hyperaldosteronism. Nature communications. 2019;10(1):4678.

39. De Sousa K, Boulkroun S, Baron S, Nanba K, Wack M, Rainey WE, et al. Genetic, Cellular, and Molecular Heterogeneity in Adrenals With Aldosterone-Producing Adenoma. Hypertension. 2020;75(4):1034–44.

40. Nivoit P, Mathivet T, Wu J, Salemkour Y, Sankar DS, Baudrie V, et al. Autophagy protein 5 controls flow-dependent endothelial functions. Cell Mol Life Sci. 2023;80(8):210.

41. Koster J, and Rahmann S. Snakemake--a scalable bioinformatics workflow engine. Bioinformatics. 2012;28(19):2520–2.

42. Chen S, Zhou Y, Chen Y, and Gu J. fastp: an ultra-fast all-in-one FASTQ preprocessor. Bioinformatics. 2018;34(17):i884–i90.

43. Dobin A, Davis CA, Schlesinger F, Drenkow J, Zaleski C, Jha S, et al. STAR: ultrafast universal RNA-seq aligner. Bioinformatics. 2013;29(1):15–21.

44. Liao Y, Smyth GK, and Shi W. featureCounts: an efficient general purpose program for assigning sequence reads to genomic features. Bioinformatics. 2014;30(7):923–30.

45. Ewels P, Magnusson M, Lundin S, and Kaller M. MultiQC: summarize analysis results for multiple tools and samples in a single report. Bioinformatics. 2016;32(19):3047–8.

46. Love MI, Huber W, and Anders S. Moderated estimation of fold change and dispersion for RNA-seq data with DESeq2. Genome Biol. 2014;15(12):550.

47. Gene Ontology C. The Gene Ontology resource: enriching a GOld mine. Nucleic Acids Res. 2021;49(D1):D325–D34.

48. Mi H, Muruganujan A, Ebert D, Huang X, and Thomas PD. PANTHER version 14: more genomes, a new PANTHER GO-slim and improvements in enrichment analysis tools. Nucleic Acids Res. 2019;47(D1):D419–D26.

49. Binns D, Dimmer E, Huntley R, Barrell D, O’Donovan C, and Apweiler R. QuickGO: a web-based tool for Gene Ontology searching. Bioinformatics. 2009;25(22):3045–6.

50. Fang Z, Liu X, and Peltz G. GSEApy: a comprehensive package for performing gene set enrichment analysis in Python. Bioinformatics. 2023;39(1).

51. Kuleshov MV, Jones MR, Rouillard AD, Fernandez NF, Duan Q, Wang Z, et al. Enrichr: a comprehensive gene set enrichment analysis web server 2016 update. Nucleic Acids Res. 2016;44(W1):W90–7.

52. Kanehisa M, and Goto S. KEGG: kyoto encyclopedia of genes and genomes. Nucleic Acids Res. 2000;28(1):27–30.

53. Cokelaer T, Pultz D, Harder LM, Serra-Musach J, and Saez-Rodriguez J. BioServices: a common Python package to access biological Web Services programmatically. Bioinformatics. 2013;29(24):3241–2.

54. Bolz SS, Vogel L, Sollinger D, Derwand R, Boer C, Pitson SM, et al. Sphingosine kinase modulates microvascular tone and myogenic responses through activation of RhoA/Rho kinase. Circulation. 2003;108(3):342–7.

55. Peter BF, Lidington D, Harada A, Bolz HJ, Vogel L, Heximer S, et al. Role of sphingosine-1-phosphate phosphohydrolase 1 in the regulation of resistance artery tone. Circulation research. 2008;103(3):315–24.

56. Kuhn R, Schwenk F, Aguet M, and Rajewsky K. Inducible gene targeting in mice. Science. 1995;269(5229):1427–9.

57. Siedlinski M, Nosalski R, Szczepaniak P, Ludwig-Galezowska AH, Mikolajczyk T, Filip M, et al. Vascular transcriptome profiling identifies Sphingosine kinase 1 as a modulator of angiotensin II-induced vascular dysfunction. Sci Rep. 2017;7:44131.

58. Pham TH, Baluk P, Xu Y, Grigorova I, Bankovich AJ, Pappu R, et al. Lymphatic endothelial cell sphingosine kinase activity is required for lymphocyte egress and lymphatic patterning. The Journal of experimental medicine. 2010;207(1):17–27.

59. Zhang J, and Crowley SD. Role of T lymphocytes in hypertension. Curr Opin Pharmacol. 2015;21:14–9.

60. Machnik A, Neuhofer W, Jantsch J, Dahlmann A, Tammela T, Machura K, et al. Macrophages regulate salt-dependent volume and blood pressure by a vascular endothelial growth factor-C-dependent buffering mechanism. Nature medicine. 2009;15(5):545–52.

61. Hallisey VM, and Schwab SR. Get me out of here: Sphingosine 1-phosphate signaling and T cell exit from tissues during an immune response. Immunol Rev. 2023.

62. Niazi H, Zoghdani N, Couty L, Leuci A, Nitzsche A, Allende ML, et al. Murine platelet production is suppressed by S1P release in the hematopoietic niche, not facilitated by blood S1P sensing. Blood Adv. 2019;3(11):1702–13.

63. Guan Z, Singletary ST, Cook AK, Hobbs JL, Pollock JS, and Inscho EW. Sphingosine-1-phosphate evokes unique segment-specific vasoconstriction of the renal microvasculature. J Am Soc Nephrol. 2014;25(8):1774–85.

64. Wang W, Shen J, Cui Y, Jiang J, Chen S, Peng J, et al. Impaired sodium excretion and salt-sensitive hypertension in corin-deficient mice. Kidney Int. 2012;82(1):26–33.

65. Kono M, Mi Y, Liu Y, Sasaki T, Allende ML, Wu YP, et al. The sphingosine-1-phosphate receptors S1P1, S1P2, and S1P3 function coordinately during embryonic angiogenesis. The Journal of biological chemistry. 2004;279(28):29367–73.

66. Akahoshi N, Ishizaki Y, Yasuda H, Murashima YL, Shinba T, Goto K, et al. Frequent spontaneous seizures followed by spatial working memory/anxiety deficits in mice lacking sphingosine 1-phosphate receptor 2. Epilepsy Behav. 2011;22(4):659–65.

67. Del Gaudio I, Rubinelli L, Sasset L, Wadsack C, Hla T, and Di Lorenzo A. Endothelial Spns2 and ApoM Regulation of Vascular Tone and Hypertension Via Sphingosine-1-Phosphate. J Am Heart Assoc. 2021;10(14):e021261.

68. Hu G, Zhu Q, Wang W, Xie D, Chen C, Li PL, et al. Collecting duct-specific knockout of sphingosine-1-phosphate receptor 1 aggravates DOCA-salt hypertension in mice. J Hypertens. 2021;39(8):1559–66.

69. Makhanova N, Sequeira-Lopez ML, Gomez RA, Kim HS, and Smithies O. Disturbed homeostasis in sodium-restricted mice heterozygous and homozygous for aldosterone synthase gene disruption. Hypertension. 2006;48(6):1151–9.

70. Kharel Y, Huang T, Salamon A, Harris TE, Santos WL, and Lynch KR. Mechanism of sphingosine 1-phosphate clearance from blood. The Biochemical journal. 2020;477(5):925–35.

71. Ye B, Bradshaw AD, Abrahante JE, Dragon JA, Haussler TN, Bell SP, et al. Left Ventricular Gene Expression in Heart Failure With Preserved Ejection Fraction-Profibrotic and Proinflammatory Pathways and Genes. Circ Heart Fail. 2023;16(8):e010395.

72. Clay H, Wilsbacher LD, Wilson SJ, Duong DN, McDonald M, Lam I, et al. Sphingosine 1-phosphate receptor-1 in cardiomyocytes is required for normal cardiac development. Dev Biol. 2016;418(1):157–65.

73. Keul P, van Borren MM, Ghanem A, Muller FU, Baartscheer A, Verkerk AO, et al. Sphingosine-1-Phosphate Receptor 1 Regulates Cardiac Function by Modulating Ca2+ Sensitivity and Na+/H+ Exchange and Mediates Protection by Ischemic Preconditioning. J Am Heart Assoc. 2016;5(5).

74. Jorgensen R, Katta M, Wolfe J, Leach DF, Lavelle B, Chun J, et al. Deletion of Sphingosine 1-Phosphate receptor 1 in cardiomyocytes during development leads to abnormal ventricular conduction and fibrosis. Physiol Rep. 2021;9(19):e15060.

75. Kono M, Tucker AE, Tran J, Bergner JB, Turner EM, and Proia RL. Sphingosine-1-phosphate receptor 1 reporter mice reveal receptor activation sites in vivo. The Journal of clinical investigation. 2014;124(5):2076–86.

76. Choi RH, Tatum SM, Symons JD, Summers SA, and Holland WL. Ceramides and other sphingolipids as drivers of cardiovascular disease. Nat Rev Cardiol. 2021;18(10):701–11.

77. Kovilakath A, Jamil M, and Cowart LA. Sphingolipids in the Heart: From Cradle to Grave. Front Endocrinol (Lausanne). 2020;11:652.

78. Yatomi Y, Igarashi Y, Yang L, Hisano N, Qi R, Asazuma N, et al. Sphingosine 1-phosphate, a bioactive sphingolipid abundantly stored in platelets, is a normal constituent of human plasma and serum. J Biochem. 1997;121(5):969–73.

79. Carey RM, Moran AE, and Whelton PK. Treatment of Hypertension: A Review. JAMA. 2022;328(18):1849–61.

80. Daugherty SL, Powers JD, Magid DJ, Tavel HM, Masoudi FA, Margolis KL, et al. Incidence and prognosis of resistant hypertension in hypertensive patients. Circulation. 2012;125(13):1635–42.

81. Schiffrin EL. How Structure, Mechanics, and Function of the Vasculature Contribute to Blood Pressure Elevation in Hypertension. Can J Cardiol. 2020;36(5):648–58.

82. Yagi K, Lidington D, Wan H, Fares JC, Meissner A, Sumiyoshi M, et al. Therapeutically Targeting Tumor Necrosis Factor-alpha/Sphingosine-1-Phosphate Signaling Corrects Myogenic Reactivity in Subarachnoid Hemorrhage. Stroke. 2015;46(8):2260–70.

83. Rawlings AM, Juraschek SP, Heiss G, Hughes T, Meyer ML, Selvin E, et al. Association of orthostatic hypotension with incident dementia, stroke, and cognitive decline. Neurology. 2018;91(8):e759–e68.

84. Weigel C, Huttner SS, Ludwig K, Krieg N, Hofmann S, Schroder NH, et al. S1P lyase inhibition protects against sepsis by promoting disease tolerance via the S1P/S1PR3 axis. EBioMedicine. 2020;58:102898.

85. Saner FH, Stueben BO, Hoyer DP, Broering DC, and Bezinover D. Use or Misuse of Albumin in Critical Ill Patients. Diseases. 2023;11(2).

86. Galvani S, and Hla T. Quality Versus Quantity: Making HDL Great Again. Arteriosclerosis, thrombosis, and vascular biology. 2017;37(6):1018–9.

87. Sattler K, Graler M, Keul P, Weske S, Reimann CM, Jindrova H, et al. Defects of High-Density Lipoproteins in Coronary Artery Disease Caused by Low Sphingosine-1-Phosphate Content: Correction by Sphingosine-1-Phosphate-Loading. J Am Coll Cardiol. 2015;66(13):1470–85.

88. Periayah MH, Halim AS, and Mat Saad AZ. Mechanism Action of Platelets and Crucial Blood Coagulation Pathways in Hemostasis. Int J Hematol Oncol Stem Cell Res. 2017;11(4):319–27.

89. Mathiesen Janiurek M, Soylu-Kucharz R, Christoffersen C, Kucharz K, and Lauritzen M. Apolipoprotein M-bound sphingosine-1-phosphate regulates blood-brain barrier paracellular permeability and transcytosis. Elife. 2019;8.

90. Yang JM, Park CS, Kim SH, Noh TW, Kim JH, Park S, et al. Dll4 Suppresses Transcytosis for Arterial Blood-Retinal Barrier Homeostasis. Circulation research. 2020;126(6):767–83.

91. Yang AC, Stevens MY, Chen MB, Lee DP, Stahli D, Gate D, et al. Physiological blood-brain transport is impaired with age by a shift in transcytosis. Nature. 2020;583(7816):425–30.

92. Means CK, and Brown JH. Sphingosine-1-phosphate receptor signalling in the heart. Cardiovascular research. 2009;82(2):193–200.

93. Ji X, Chen Z, Wang Q, Li B, Wei Y, Li Y, et al. Sphingolipid metabolism controls mammalian heart regeneration. Cell metabolism. 2024;36(4):839–56 e8.

94. Xie T, Chen C, Peng Z, Brown BC, Reisz JA, Xu P, et al. Erythrocyte Metabolic Reprogramming by Sphingosine 1-Phosphate in Chronic Kidney Disease and Therapies. Circulation research. 2020;127(3):360–75.

95. Sun K, Zhang Y, D’Alessandro A, Nemkov T, Song A, Wu H, et al. Sphingosine-1-phosphate promotes erythrocyte glycolysis and oxygen release for adaptation to high-altitude hypoxia. Nature communications. 2016;7:12086.

96. Thomas N, Schroder NH, Nowak MK, Wollnitzke P, Ghaderi S, von Wnuck Lipinski K, et al. Sphingosine-1-phosphate suppresses GLUT activity through PP2A and counteracts hyperglycemia in diabetic red blood cells. Nature communications. 2023;14(1):8329.

